# Neural evidence that humans reuse strategies to solve new tasks

**DOI:** 10.1101/2024.06.10.598294

**Authors:** Sam Hall-McMaster, Momchil S. Tomov, Samuel J. Gershman, Nicolas W. Schuck

## Abstract

Generalisation from past experience is an important feature of intelligent systems. When faced with a new task, one efficient computational approach is to evaluate solutions to earlier tasks as candidates for reuse. Consistent with this idea, we found that human participants (n=38) learned optimal solutions to a set of training tasks and generalised them to novel test tasks in a reward selective manner. This behaviour was consistent with a computational process based on the successor representation known as successor features and generalised policy improvement (SF&GPI). Neither model-free perseveration or model-based control using a complete model of the environment could explain choice behaviour. Decoding from functional magnetic resonance imaging data revealed that solutions from the SF&GPI algorithm were activated on test tasks in visual and prefrontal cortex. This activation had a functional connection to behaviour in that stronger activation of SF&GPI solutions in visual areas was associated with increased behavioural reuse. These findings point to a possible neural implementation of an adaptive algorithm for generalisation across tasks.

## Introduction

The ability to flexibly generalise from past experience to new situations is central to intelligence. Humans often excel at this (Dekker et al., 2022; Luettgau et al., 2023; Xia & Collins, 2021) and the reasons have long intrigued cognitive neuroscientists. A recurring theme is that neural systems should store structural information extracted from past experiences (Behrens et al., 2018; Tervo et al., 2016). Structural information specifies the *form* shared by solutions to a given class of problems, abstracting over the *content* that is specific to each problem. When new situations arise, this information can be retrieved and combined with situation-specific content to make decisions.

Although this approach is maximally flexible, humans often use simpler forms of generalisation (Carvalho et al., 2024). Recent data have shown that humans tend to solve new tasks by reusing solutions from earlier tasks (Tomov et al., 2021). This relies on an algorithm known as “generalised policy improvement” (GPI; Barreto et al. 2017, 2018, 2020). GPI achieves efficient generalisation by storing a set of solutions (policies) that can be evaluated and selected for reuse. Importantly, a feature-based generalisation of the successor representation (“successor features” or SFs) can be harnessed to support the identification of the optimal policy among those stored in memory. The resulting algorithm, SF&GPI, is able to efficiently solve new tasks (Barreto et al. 2017, 2018, 2020) and predict human generalisation behaviour (Tomov et al., 2021), when tasks are situated in a common environment and each task is associated with a distinct reward function.

Here we investigated whether the human brain implements this flexible and efficient form of generalisation. If people generalise their past experiences using SF&GPI, it should be possible to detect its components in their brain activity. We developed three neural predictions based on this premise. First, we predicted that successful past policies would be represented in brain activity when people are exposed to new tasks. Second, we predicted that these policies would be prioritised, showing stronger activation than unsuccessful past policies. Third, we predicted a corresponding representation of the features associated with successful past policies, as these are used in the model to compute expected rewards.

Past research from cognitive neuroscience provides important clues about where these predictions should be observed. The dorsolateral prefrontal cortex (DLPFC) has been proposed as a region that encodes policies (Botvinick & An, 2008; Fine & Hayden, 2022) and supports context-dependent action (Badre & Nee, 2017; Flesch et al., 2022; Frith, 2000; Jackson et al., 2021; Rowe et al. 2000). Based on this literature, DLPFC is a candidate region in which the encoding of successful past policies could be detected when people are exposed to new situations. The medial temporal lobe (MTL) and orbitofrontal cortex (OFC) have been proposed as regions that encode predictive representations about future states (De Cothi & Barry, 2020; Geerts et al. 2020; Muhle-Karbe et al., 2023; Stachenfeld et al. 2017; Wimmer & Büchel, 2019). Based on this literature, MTL and OFC are candidate regions in which features associated with successful past policies might be detected.

To test these predictions, participants completed a multi-task learning experiment during functional magnetic resonance imaging (fMRI). The experiment included training tasks that participants could use to learn about their environment, and test tasks to probe their generalisation strategy. Different reward functions were used to define different tasks in the experiment. To summarise the main results, we found that participants learned optimal solutions (policies) to the training tasks, and generalised them to test tasks in a reward selective manner. Participant choices at test were more similar to an SF&GPI algorithm than an MB algorithm with full knowledge of the environment. Participant choices were also distinct from a model-free process. Neural results showed that optimal solutions from the training tasks could be decoded above chance during test tasks in DLPFC and (surprisingly) in occipitotemporal cortex (OTC). These solutions were also prioritised. Decoding evidence for the optimal training solutions at test was higher than alternative solutions that promised larger rewards. These results provide new insights into how a sophisticated policy reuse algorithm might be implemented in the brain.

## Results

To test whether SF&GPI computations were evident in brain activity, participants completed a gem collector game inside an MRI scanner (Fig. 1). The cover story was that a criminal mastermind had hidden rare gems in different cities around the world. Participants needed to retrieve the gems to sell them for as much profit as possible. The first screen on each trial showed a set of market values that indicated the profits or losses associated with reselling each gem. Market prices could range from $2 per gem to -$2 per gem. After seeing the market values, participants faced a choice between four cities (Sydney, Tokyo, New York, London) shown in a random order on screen. Each city contained the same three gem stones (triangle, square, circle), but in different amounts. The profit earned for a particular choice was based on the combination of market values and the recovered gems. The selling price of each gem stone was first multiplied by its respective number in the chosen city, and rewards were then summed across gem stones to arrive at the total profit. Embedded in this trial structure were two important abstract elements. First, market values were reward functions that defined the task participants needed to solve on a given trial. If triangular gems had a high market value for example, and all other factors were equal, participants would need to locate the city with the most triangular gems. Second, the gem numbers in each city defined the state features for that city, information that could be reused to guide decisions on new tasks.

**Figure 1.**
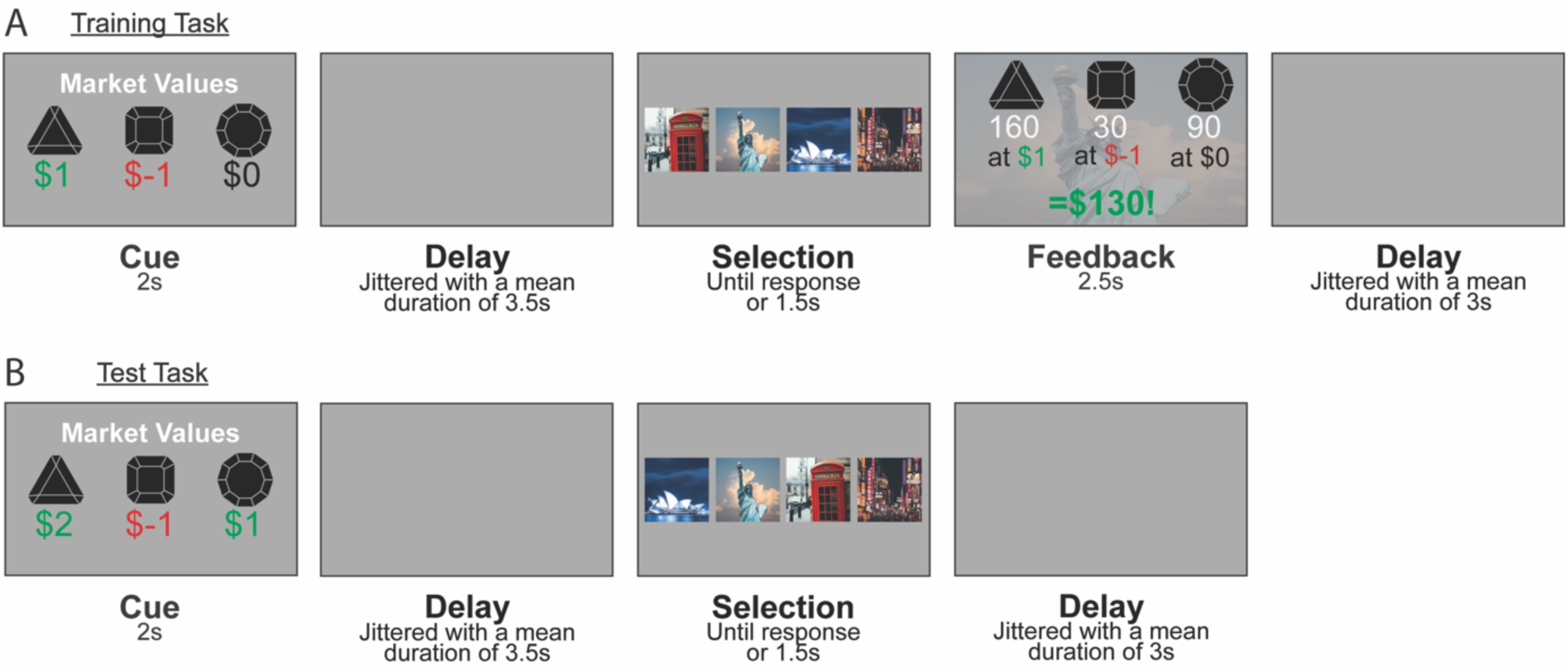
Experimental design. Participants competed a gem collector game while their brain activity was measured with fMRI. On each trial in the experiment, gems with distinct shapes could be resold for either a gain or a loss. Participants made a choice between four cities from around the world, each leading to a distinct collection of gems. To maximise profit overall, participants needed to choose the city best suited to the selling prices shown on each trial. Each block consisted of 32 training trials that included feedback (shown in A), followed by a mixture of 16 training trials with feedback and 20 test trials without feedback (shown in B). The gem collection associated with each city changed from block to block. **A**: An example training task. Following presentation of the market values, participants selected a city and saw a feedback screen. The feedback screen revealed the gem numbers in the selected city and the profit earned on the current trial. Four training tasks were used in the experiment and were designed so that two of the four cities were optimal across training tasks. **B**: An example test task. On test tasks, participants saw a set of market values that had not appeared during training and selected a city but did not see the outcome of their decision. Four test tasks were used in the experiment and were designed so that the two cities that were previously suboptimal now offered the highest returns. A model-based agent with a complete model of the environment is expected to be sensitive to this change. A “memory-based” SF&GPI agent that evaluates earlier task solutions as candidates for generalisation is expected to choose among the cities that were optimal during training.

Trials were ordered in a specific way to test the SF&GPI model. When beginning each block, participants did not know how many gems were present in each city and needed to learn this information by making decisions and observing the outcomes. This was possible during the first 32 trials of the block, which included feedback after each choice (Fig 1A). We refer to trials with feedback as *training trials* hereafter. Across training trials, participants encountered four training tasks that each had a unique reward function (i.e. a unique market value cue shown at the beginning of the trial). Two training tasks resulted in high rewards when city A was selected (e.g. Sydney), while the alternative cities resulted in losses or marginal reward. The other two training tasks resulted in high rewards when city B was selected (e.g. Tokyo), but losses or marginal reward when the alternative cities were selected. This meant that in effect participants needed to learn two optimal policies to perform well on training trials. One optimal policy could be used for two training tasks and another could be used for the remaining two training tasks.

Following the initial learning phase, participants were given 20 *test trials* (Fig. 1B) interspersed with 16 training trials. The test trials differed from training trials in that they introduced four new tasks (market value cues) and did not provide feedback after each choice. The test trials were designed so that the two cities resulting in losses or marginal reward during training (e.g. New York and London) were now the most rewarding. This experimental setup allowed us to dissociate two main generalisation strategies. A model-based agent with full knowledge of the environment would compute expected rewards for all four cities and choose the one that was objectively most rewarding. A model-based agent would therefore enact different policies on the test tasks compared to the training tasks. In contrast, an SF&GPI agent that stores and evaluates the best cities from training would compute expected rewards based on the optimal training policies. An SF&GPI agent would therefore choose the more rewarding city among the optimal training policies for each test task, but would not enact polices that had been unrewarding during training.

Participants completed six blocks of 68 trials in total. Training tasks had the following market values: w_train_ = {[1, −1, 0], [−1, 1, 0], [1, −2, 0], [−2, 1, 0]}. Test tasks had the following market values: w_test_ = {[2, -1, -1], [-1, 1, 1], [1, -1, 1], [1, 1, -1]}. Gem numbers (state features) had the following values: ɸ = {[120, 50, 110], [90, 80, 190], [140, 150, 40], [60, 200, 20]}. Reward for each combination of w and ɸ is presented in the methods (Fig. 4). The vector elements in each triplet were shuffled using a shared rule before a new block (e.g. all vectors were reordered to [1,3,2]). The mapping between cities and features (gem triplets) was also changed. These changes created the appearance of new training and test tasks in each block while preserving the structure of the experiment, in which two of the four options always proved to be best in training trials but suboptimal in test trials. Performance was incentivised with a monetary bonus that was based on the total profit accrued over all trials in the scanner game. Before the experiment took place, participants were informed about how market values and gem numbers were combined to calculate the profit on each trial. Participants also completed 80 training trials with different state features and a different task theme in preparation for the session, and 20 training trials with different state features prior to scanning.

### Participants learned optimal policies for training tasks

We first examined performance on the training tasks (Fig. 2A-E). This included training trials from the initial learning phase and training trials that were interspersed between test trials. The average reward earned per choice was higher than what would be expected if participants were choosing at random (M=49.77, SD=10.61, M_random_=-54.38, SD_random_=6.16, *t*(74)=52.34, corrected *p*<0.001) suggesting successful learning. To understand training performance in more detail, we examined which choice options were selected. Training tasks were designed so that the most rewarding choice on each trial would lead to gem triplet ɸ(1) or gem triplet ɸ(4). Gem triplets are called feature triplets hereafter. Consistent with the training structure, cities associated with ɸ(1) and ɸ(4) were chosen significantly more often than the other cities (Fig. 2C). ɸ(1) was reached more often than ɸ(2) or ɸ(3) (M_ɸ(1)_=41.66% of training trials, SD_ɸ(1)_=3.44, M_ɸ(2)_=6.52%, SD _ɸ(2)_=2.70, *t*(37)=37.78, corrected *p*<0.001; M_ɸ(3)_=6.38%, SD_ɸ(3)_=2.71, *t*(37)=37.95, corrected *p*<0.001). ɸ(4) was similarly reached more often than ɸ(2) or ɸ(3) (M_ɸ(4)_=45.28% of training trials, SD_ɸ(4)_=2.02, *t*_ɸ(*4) vs.* ɸ(*2)*_(37)=45.98, corrected *p*<0.001; *t*_ɸ(*4) vs.* ɸ(*3)*_(37)=45.42, corrected *p*<0.001). Choices leading to ɸ(1) and ɸ(4) were more comparable in number. However, ɸ(4) was reached more often than ɸ(1) (*t*(37)=5.34, corrected *p*<0.001). To understand the optimality of these decision patterns, we examined how often participants made the optimal choice on training trials. The percentage of optimal choices was significantly above chance (M=82.94%, SD=6.36, chance=25%, *t*(37)=56.16, corrected *p*<0.001, Fig. 2E) indicating that participants acquired the optimal training policies. Together, these results indicate that participants acquired robust and effective decision strategies to maximise reward on the training tasks.

**Figure 2.**
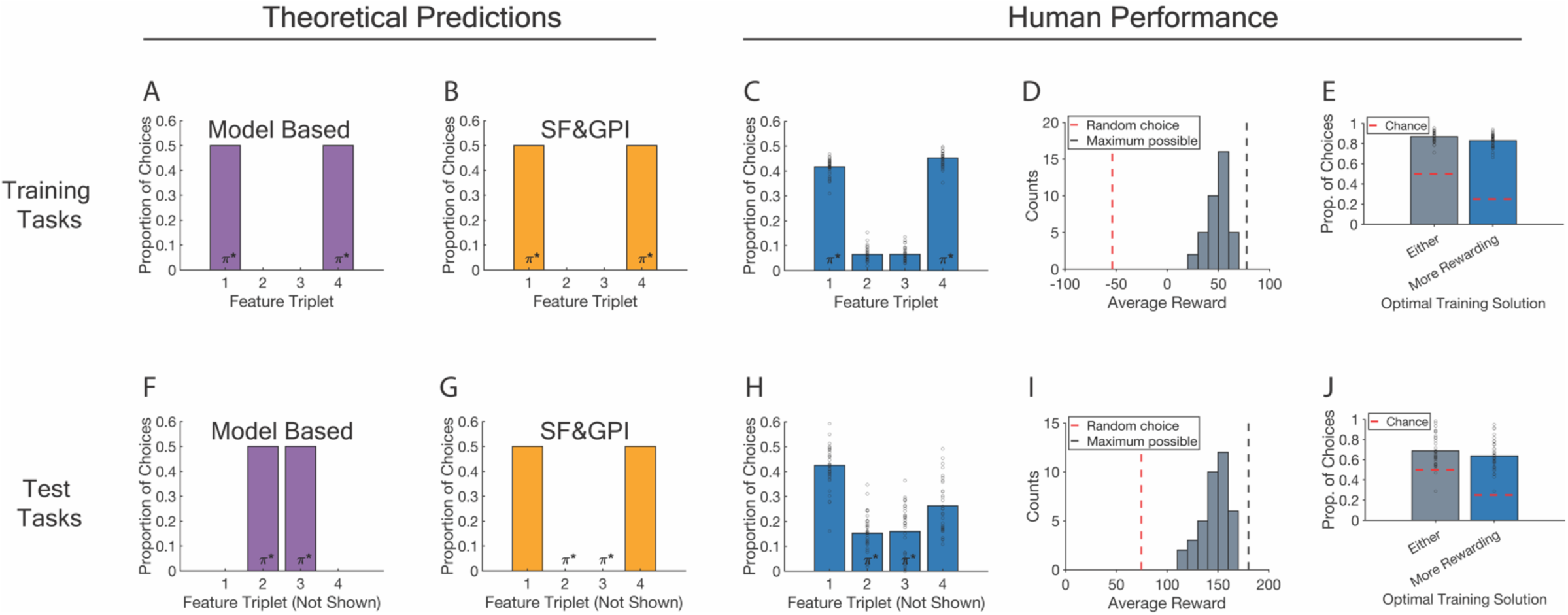
Behavioural results. Columns are grouped into theoretical predictions and human performance. Rows are grouped into training and test tasks. **A, B, F, G**: Theoretical predictions are shown for a model-based algorithm on training (**A**) and test tasks (**F**), as well as an SF&GPI algorithm on training (**B**) and test tasks (**G**). Each theoretical plot shows the predicted proportion of choices leading to each feature (gem) triplet. Theoretical plots show expected choice profiles after model convergence on training tasks and do not include choices during learning. **C, H**: Human choice profiles for training (**C**) and test tasks (**H**). The axis structure matches the theoretical plots. **D, I:** Performance histograms. The x-axis shows the average reward earned per trial within 10-point bins, and the y-axis shows number of participants with this average. The dashed red line indicates performance expected from random choices. The dashed grey line indicates the maximum performance possible. **E-J:** Use of the optimal training solutions (policies) on training tasks (**E**) and their reuse on test tasks (**J**). Both plots show the proportion of choices in which either optimal training policy was used (left bars) and the proportion of choices in which the more rewarding one was used (right bars). Dashed red lines indicate chance. **A, B, C, F, G, H:** π* denotes feature triplets associated with optimal training and test policies. Optimal policies for training tasks lead to feature triplet ɸ(1) or ɸ(4) and optimal policies for test tasks lead to ɸ(2) or ɸ(3). **C, H, E, J:** Dots show data points from individual participants. **A-J:** Materials to reproduce these panels are available at https://gin.g-node.org/sam.hall-mcmaster/sfgpi-behavioural-analysis.

### Participants transferred optimal training policies to test tasks

Having shown that participants learned the optimal training policies, we turned our attention to the test tasks (Fig. 2F-J). The test tasks were designed so that choices leading to feature triplets ɸ(2) or ɸ(3) were the most rewarding. A model-based agent was expected to compute anticipated rewards under all available policies and therefore make different choices on test tasks compared to the training tasks. An SF&GPI agent that stores information about the best cities from training was expected to compute anticipated rewards only under the optimal training policies. This would result in continued choices to reach ɸ(1) and ɸ(4), rather than switching to ɸ(2) and ɸ(3). Although decisions for ɸ(1) and ɸ(4) were suboptimal, we expected that participants would still select the option that generated higher reward among this set.

Consistent with an SF&GPI algorithm, participants continued using optimal policies from training on most test trials (M=68.81%, SD=15.58, chance=50%, *t*(37)=7.44, *p*<0.001, Fig. 2J). The choice profile on test trials was also similar to the profile seen on training trials (Fig. 2H). The city associated with feature triplet ɸ(1) was selected significantly more often than cities that led to ɸ(2) and ɸ(3) (M_ɸ(1)_=42.52% of test trials, SD_ɸ(1)_=8.27, M_ɸ(2)_=15.23%, SD_ɸ(2)_=7.43, *t*(37)=11.48, corrected *p*<0.001; M_ɸ(3)_=15.96%, SD_ɸ(3)_=10.23, *t*(37)=9.78, corrected *p*<0.001). The city associated with feature triplet ɸ(4) was similarly selected significantly more often than those associated with ɸ(2) and ɸ(3) (M_ɸ(4)_=26.29% of test trials, SD_ɸ(4)_=10.64, *t*_ɸ(*4) vs.* ɸ(*2)*_(37)=4.14, corrected *p*<0.001; *t*_ɸ(*4) vs.* ɸ(*3)*_(37)=3.17, corrected *p*=0.006). ɸ(1) was reached more often than ɸ(4) during test tasks (*t*(37)=9.12, corrected *p*<0.001). While most choices were technically suboptimal, participants still performed well on test tasks overall. The average reward per choice was significantly higher than the reward expected from random choice (M=147.00, SD=13.94, M_random_=76.89, SD_random_=7.84, *t*(74)= 27.02, *p*<0.001, Fig. 2I). This was due to participants selecting the more rewarding solution among the optimal training policies on test tasks significantly more often than chance (M=63.71%, SD=14.22, chance=0.25, *t*(37)=16.78, corrected *p*<0.001, Fig. 2J). On trials where participants used either of the two optimal training policies, the more rewarding one was indeed selected in most cases (M=92.86%, SD=5.21, chance=0.5, *t*(37)=50.74, corrected *p*<0.001). The proportion of SF&GPI-predicted choices on test tasks was positively related to the proportion of optimal choices during training but not to a significant degree (Spearman’s *Rho*=0.294, corrected *p*=0.073).

Auxiliary behavioural results are presented in the supplementary information. Training performance reached an asymptote before test tasks were introduced (Fig. S1) and participants could be separated into subgroups based on how closely their behaviour matched the SF&GPI predictions (Fig. S2). The SF&GPI-predicted choice was made significantly more often than the alternatives on 3 of the 4 individual test tasks (Fig. S3), with the remaining test task showing an even split between SF&GPI and MB choice, an interesting behavioural signature of Universal Value Function Approximation (Fig. S4).

Generalisation performance could not be explained by a simple (model-free) perseveration process (Fig. S5). The experiment also included an assessment of participants’ explicit knowledge. Following scanning, participants estimated the number of gems that were present in each city during the final block of the experiment (Fig. S6). While we did not detect a difference in the mean gem estimate error across policies (corrected *p*-values>0.09, Fig. S6A), we observed a significant positive correlation between the estimate error for cities that were suboptimal during training and SF&GPI-consistent choices on test trials (Spearman’s *Rho*=0.493, corrected *p*=0.016, Fig. S6B). This suggests a possible link between noisy memory for feature triplets associated with the suboptimal training policies and the reuse of successful past solutions.

Taken together, the results in this section indicate that participants were choosing among the optimal training policies on test trials in a reward-sensitive manner, a behavioural profile more consistent with the predictions of an SF&GPI algorithm than an MB algorithm operating on a full model of the environment.

### Optimal training policies could be decoded during test tasks

Having established that behaviour was consistent with the predictions of an SF&GPI model, we tested its neural predictions on test tasks: 1) that optimal training policies would be activated as decision candidates; 2) that their activation strength would be higher than alternative policies; and 3) that features associated with the optimal training policies would also be represented. We tested these predictions in four brain regions. Predictions 1-2 were expected in DLPFC due to its proposed role in policy encoding (Botvinick & An, 2008; Fine & Hayden, 2022) and context-dependent action (Badre & Nee, 2017; Flesch et al., 2022; Frith, 2000; Jackson et al., 2021; Rowe et al. 2000). Prediction 3 was expected in MTL and OFC due to research implicating these regions in coding predictive information about future states (De Cothi & Barry, 2020; Geerts et al. 2020; Muhle-Karbe et al., 2023; Stachenfeld et al. 2017; Wimmer & Büchel, 2019). OTC was examined as a final region due to its central role in early fMRI decoding studies and its continued inclusion in recent ones (Haxby et al., 2001; Muhle-Karbe et al., 2023; Wittkuhn et al., 2021, 2024). We first focus on policy activation (predictions 1-2).

To examine policy activation, decoders based on logistic regression were trained to distinguish the four cities seen during feedback on the training tasks (i.e. the cities served as training labels). One measurement volume (TR) per eligible trial was used as input for decoder training, taken 4-6s after feedback onset to account for the hemodynamic delay. Decoders were trained separately for each region of interest (ROI). The specific time lag used for each ROI was determined in validation analyses of the training tasks, independent from our predictions about test task activity (see Fig. S7). Validation analyses further showed that decoder training locked to feedback onset on training tasks was more effective than decoder training locked to task cue onset (see Fig. S8). The decoders were applied to each TR on held out test trials in a leave-one-run-out cross validation procedure. This resulted in a decoding probability time course for each city on each test trial, which reflected the evidence that a particular city stimulus was encoded in the fMRI signal. To account for class imbalances in the decoder training set, we repeated the analysis 100 times using random subsampling that matched the number of trials per category (M=53 trials per training class, SD=6.48, minimum=37, maximum=65). Decoding probabilities based on the cities were then coded into the following categories: 1) the more rewarding policy (among the optimal training policies); 2) the less rewarding policy (among the optimal training policies); 3) the objective best policy; and 4) the remaining policy.

We first tested the prediction that optimal training policies would be activated on test tasks, by comparing decoding evidence for the optimal training policies against chance in each ROI (Fig. 3B). Test trials lasted 10s on average (Fig 1B). Decoding evidence was therefore averaged from +2.5s to +12.5s following test trial onset to account for the hemodynamic delay. This revealed significant average decoding evidence for the more rewarding training policy during test tasks in OTC and DLPFC (OTC: M=28.07%, SD=2.17, *t*(37)=8.57, corrected *p*<0.001; DLPFC: M=25.85%, SD=1.51, *t*(37)=3.40, corrected *p*=0.011), but not in MTL or OFC (MTL: M=25.17%, SD=1.43, *t*(37)=0.72, corrected *p*=0.744; OFC: M=25.40%, SD=1.37, *t*(37)=1.79, corrected *p*=0.405). Numerical evidence for the less rewarding training policy was also detected in OTC, but this did not survive correction (M=25.97%, SD=2.34, *t*(37)=2.53, uncorrected *p*=0.016, Bonferroni-Holm corrected *p*=0.094, see the section entitled *Reactivation and stimulus processing contributed to the decoding effects* for evidence from cluster-based permutation tests). Average decoding evidence for the less rewarding training policy was not detected in the remaining ROIs (MTL: M=25.32%, SD=1.76, *t*(37)=1.12, corrected *p*=0.744; OFC: M=25.31%, SD=1.40, *t*(37)=1.38, corrected *p*=0.708; DLPFC: M=25.30%, SD=1.53, *t*(37)=1.17, corrected *p*=0.744). Decoding evidence on individual test tasks is included in the supplement (Fig. S9). These results indicate that the more rewarding among the optimal training policies was activated in OTC and DLPFC during test tasks.

**Figure 3.**
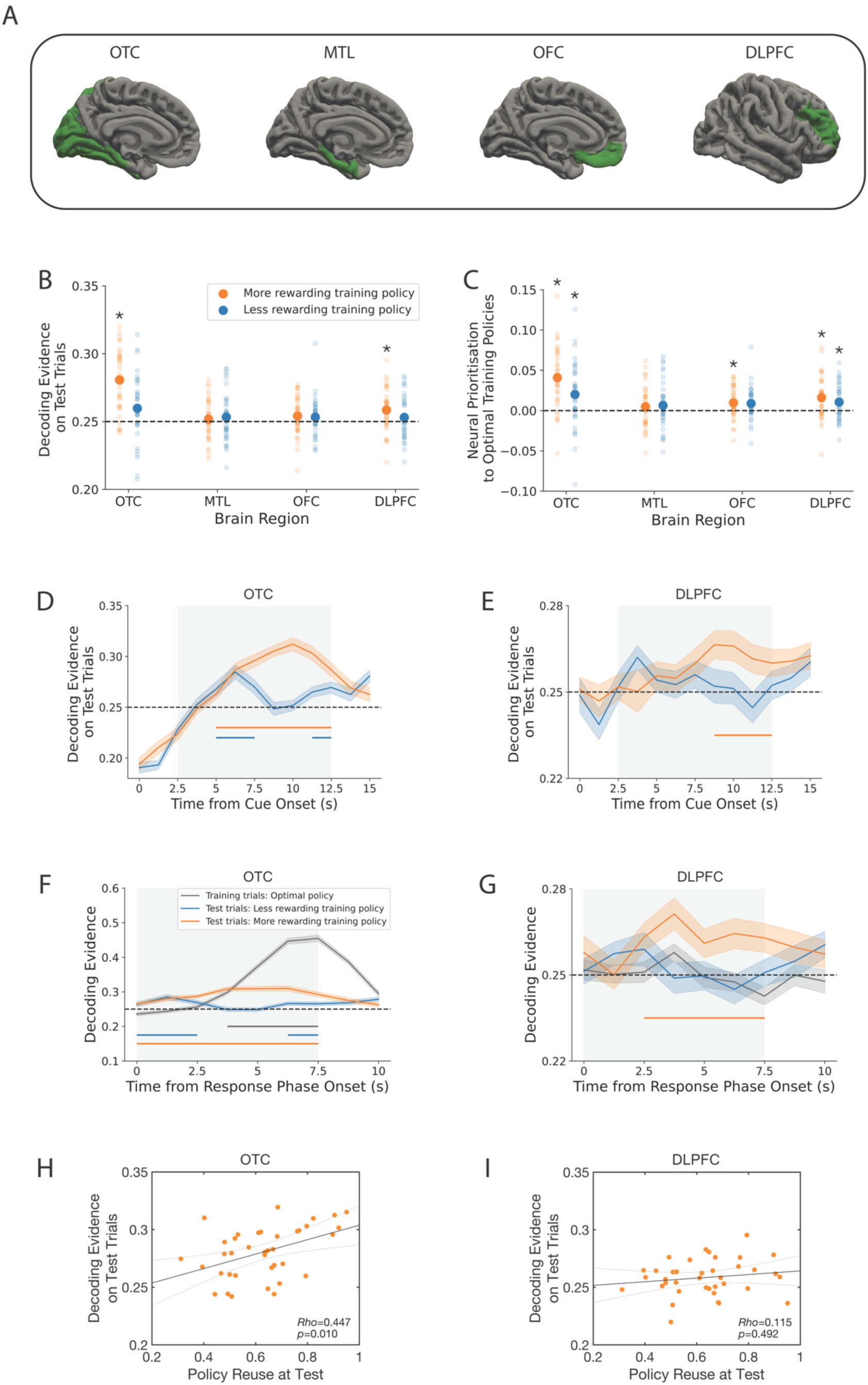
Decoding results. **A:** Brain regions of interest. The four regions include occipitotemporal cortex (OTC), the medial temporal lobe (MTL), orbitofrontal cortex (OFC) and dorsolateral prefrontal cortex (DLPFC). Regions were defined using FreeSurfer (Fischl, 2012). **B**: Decoding evidence for the optimal training policies on test tasks (y-axis) shown for each brain region (x-axis). The dashed line indicates chance. **C**: Neural prioritisation to the optimal training policies. This panel shows the difference in decoding evidence on test tasks between the optimal training policies and the objective best policy. Higher values indicate stronger evidence for the optimal training policies within the neural signal. **D-E**: Time resolved decoding evidence for the optimal training policies on test trials. Panels begin at the onset of the test cue. The initial period shows negative decoding evidence due to a control procedure. To avoid activity from the previous trial biasing our assessment of the current trial, we excluded trials in which the current policy category was selected on the previous trial. **B-E**: The effects with consistent numerical and statistical evidence across panels B-E are the activation and prioritisation of the optimal training policies in OTC, as well as the activation and prioritisation of the more rewarding training policy in DLPFC. The remaining effects in B-E did not show consistent evidence across analyses and are not interpreted further. **F-G**: Time resolved decoding evidence locked to the response phase onset. Evidence is presented for both the optimal policy on training trials and the optimal training policies on test trials. **D-G:** Coloured bars below each line indicate significant decoding clusters. Shaded error bars show the standard error of mean. **H-I**: Relationships between decoding evidence for the more rewarding training policy in a specific ROI (y-axis) and the proportion of test trials in which participants reused that policy (x-axis). Black lines indicate linear fits to the data and grey lines indicate 95% confidence intervals of the fits. **B-I**: Throughout the figure, decoding evidence for the more rewarding among the two optimal training policies is shown in orange, and the less rewarding among the two optimal training policies is shown in blue. The policy within this set that is more or less rewarding varies across test trials depending on the specific cue. **B-G:** Materials to reproduce these panels are available at https://gin.g-node.org/sam.hall-mcmaster/sfgpi-neural-analysis. **H-I:** Materials to reproduce these panels are available at https://gin.g-node.org/sam.hall-mcmaster/sfgpi-behavioural-analysis.

### Optimal training policies were prioritised on test tasks

Having established that the optimal training policies were activated on test tasks (prediction 1), we next tested whether their activation strength was higher than the other policies (prediction 2, Fig. 3C). Average decoding evidence during the test tasks was significantly higher for the more rewarding training policy than the objective best policy in OTC, OFC and DLPFC (OTC: M_diff_=4.09%, SD_diff_=3.69, *t*(37)=6.75, corrected *p*<0.001; MTL: M_diff_=0.48%, SD_diff_=2.46, *t*(38)=1.17, corrected *p*=0.743; OFC: M_diff_=0.97%, SD_diff_=1.87, *t*(38)=3.14, corrected *p*=0.030; DLPFC: M_diff_=1.61%, SD_diff_=2.59, *t*(38)=3.77, corrected *p*=0.006). Similar results were observed for the less rewarding training policy. Average decoding evidence for the less rewarding training policy was significantly higher than the objective best policy in OTC and DLPFC (OTC: M_diff_=2.00%, SD_diff_=3.95, *t*(37)=3.07, corrected *p*=0.032; DLPFC: M_diff_=1.05%, SD_diff_=2.15, *t*(37)=2.99, corrected *p*=0.035; MTL: M_diff_=0.63%, SD_diff_=2.80, *t*(37)=1.36, corrected *p*=0.722; OFC: M_diff_=0.88%, SD_diff_=1.97, *t*(37)=2.71, corrected *p*=0.060). Relative decoding scores above take the form: (evidence for training policy X) – (evidence for the objective best policy) and measure the extent to which optimal training policies were prioritised in neural activity. We found that relative decoding scores were significantly higher for the more rewarding training policy than the less rewarding one in OTC (M_diff_=2.09%, SD_diff_=2.37, *t*(37)=5.38, corrected *p*<0.001) but comparable in other ROIs (MTL: M_diff_=-0.15%, SD_diff_=1.61, *t*(37)=-0.58, corrected *p*>0.99; OFC: M_diff_=0.09%, SD_diff_=1.93, *t*(37)=0.28, corrected *p*>0.99; DLPFC: M_diff_=0.55%, SD_diff_=1.94, *t*(37)=1.73, corrected *p*=0.092). The results from testing predictions 1 and 2 suggest that successful training policies were activated and prioritised on test tasks in OTC. The more rewarding training policy was activated and prioritised in DLPFC (Fig. 3B-E). Remaining effects did not show consistent evidence when testing both predictions and are not considered further.

### Reactivation and stimulus processing contributed to the decoding effects

We next used cluster-based permutation testing (Maris & Oostenveld, 2007) to estimate when the optimal training policies were activated in the decoding time courses (window tested=2.5-12.5s post cue; 10,000 permutations). These tests revealed significant decoding evidence for the more rewarding training policy from 5-12.5s on test trials in OTC (corrected *p*<0.001) and from 8.75-12.5s in DLPFC (corrected *p*<0.001). Cluster-based permutation testing is potentially more sensitive than testing average evidence because activation that occurs in smaller subsets of time points can be identified. Consistent with this idea, cluster tests identified significant decoding evidence for the less rewarding training policy in OTC from 5-7.5s (corrected *p*<0.001) and 11.25-12.5s (corrected *p*=0.016), as well as a candidate cluster at 3.75s in DLPFC (corrected *p*=0.072). No signfiicant decoding clusters were detected for OFC or MTL (all corrected *p*-values>0.301). These results suggest that the more rewarding training policy was activated in the MRI signal from OTC about 5s after trial onset and in the signal from DLPFC about 8.75 s after trial onset. The OTC signal also had transient information about the less rewarding past policy from about 5s (Fig. 3D-E).

To assess whether these effects were impacted by the choice made on each test trial, we re-ran the cluster tests above but excluded evidence from trials where the policy category matched the choice participants made. A significant decoding cluster for the more rewarding training policy was detected in OTC from 5s-7.5s (corrected *p*=0.003) but no significant clusters were detected in DLPFC (candidate cluster at 5s, corrected *p*=0.254). The same results were seen for the less rewarding training policy. Significant decoding clusters were observed in OTC from 5s-7.5s (corrected *p*=0.003) and from 11.25s-12.5s (corrected *p*=0.021), but not in DLPFC (no candidate clusters). These results suggest that information about successful past policies could still be decoded from OTC when controlling for the choice made. This was not the case for DLPFC.

So far we have seen that OTC activated the optimal training policies and that DLPFC activated the more rewarding of the optimal training policies. These effects could be due to the neural reactivation of the policies from memory following the test cue, selective attention to specific policies when the response screen was shown, or both. To arbitrate between these possibilities, we examined decoding evidence locked to the response phase, and asked whether activation of the optimal training policies arose earlier than could be expected from the response phase alone (Fig. 3F-G). The procedure for training the decoder remained the same as the previous analyses. However, the evaluation was conducted at each time point from 0s to 7.5s relative to response phase onset rather than the cue onset. To establish the expected lag of policy decoding after the response phase onset, we examined decoding evidence on training trials using cluster-based permutation tests (window tested=0-7.5s, 10,000 permutations). Using these trials as a baseline revealed a significant decoding cluster for the optimal policy in OTC that was first detected 3.75s after the start of the response phase during training trials (significant cluster window=3.75-7.5s, corrected *p*<0.001, Fig. 3F). In contrast to this expected lag, the cluster onset was shifted earlier on test trials, with significant information about the optimal training policies present in OTC already 0s from the response phase onset (more rewarding training policy: significant cluster window=0-7.5s, corrected *p*<0.001; less rewarding past policy: first significant cluster window=0-2.5s, corrected *p*=0.001, second significant cluster window=6.25-7.5s, corrected p=0.014). In line with these results, decoding evidence in OTC at 0s from the response phase onset was significantly higher on test trials compared to training trials (more rewarding training policy: M_diff_=2.94%, SD_diff_=3.38, *t*(37)=5.29, corrected *p*<0.001; less rewarding training policy: M_diff_=2.81%, SD_diff_=3.97, *t*(37)=4.31, corrected *p*<0.001). No differences at 0s were detected for DLPFC (corrected *p* values>0.623, Fig. 3G). To summarise, information about the optimal training policies was present from response phase onset in OTC during test trials. Due to the hemodynamic delay, this implies some reactivation of the optimal training policies in OTC before the response screen was displayed. Attention to specific stimuli shown on the response screen would have then plausibly contributed to the decoding effects in OTC from around 2.5-3.75s after its onset (Fig. 3F).

### Policy activation in OTC was associated with test choices

The neural results presented above suggest that OTC and DLPFC represented information predicted under an SF&GPI algorithm. To examine whether neural coding in OTC and DLPFC had a functional connection to participant choices, we correlated average decoding strength for the more rewarding training policy (the orange dots in Fig. 3B) with the proportion of test trials in which participants generalised the more rewarding training policy. This revealed a significant positive correlation between neural activation of the more rewarding training policy in OTC and the implementation of that policy at test (Spearman’s *Rho*=0.447, corrected *p*=0.010, Fig. 3H). The equivalent correlation was not detected for DLPFC (Spearman’s *Rho*=0.115, corrected *p*=0.492, Fig. 3I). The correlation results held when using neural prioritisation towards the more rewarding training policy (the orange dots in Fig. 3C) as the neural variable (OTC: Spearman’s *Rho*=0.433, corrected *p*=0.013; DLPFC: Spearman’s *Rho*=0.161, corrected *p*=0.335). Neural coding in OTC was also directionally specific.

Increased neural evidence for the more rewarding training policy in OTC showed a significant negative relation to the proportion of optimal MB choices at test (OTC: Spearman’s *Rho*=-0.375, corrected *p*=0.041; DLPFC: Spearman’s *Rho*=-0.098, corrected *p*=0.560). These results suggest that policy coding within the OTC was associated with test choices.

### Features could not be decoded on test tasks

Having seen that optimal training policies were activated on test tasks (prediction 1) and that the evidence for them was higher than the objective best policy (prediction 2), we tested whether features associated with the optimal policies were also represented (prediction 3). This required a different decoding approach for two reasons. The first was that feature triplets ɸ(1) and ɸ(4) were consistently associated with the optimal training policies and were thus selected on most trials. Training a feature decoder on these data directly would result in large imbalances in the number of trials per class. The second reason was that the feature triplets were correlated with the reward. Feature triplets ɸ(1) and ɸ(4) often resulted in a profit and feature triplets ɸ(2) and ɸ(3) often resulted in a loss.

To circumvent these issues, we trained feature decoders on fMRI data from a separate associative memory paradigm (see *Session One* in the methods section and Fig. S10). Participants were first pre-trained on associations between visual cues and target stimuli. The target stimuli were feature numbers or cities that would later be used in the gem collector game. On each scanning trial, participants were shown a visual cue and needed to select its associated target from a choice array. Trials in which the correct number target was selected were used to train logistic decoders that could distinguish neural responses for the 12 number targets. The training process was similar to the process used for policy decoding. One measurement volume (TR) per eligible trial was used as training input, taken 4-6s after the response screen onset. The time shift and smoothing used for each ROI were the same as those used to decode policies. This approach could successfully recover the 12 number targets when tested in held-out data from the associative memory paradigm in all ROIs except MTL (Fig. S11). The trained decoders were then shown neural data from the test trials in the gem collector paradigm. This returned a cross-validated decoding probability for each feature number at each TR on each test trial. To ensure the decoding probabilities were stable, we repeated the procedure 100 times using random subsampling to match trial numbers (M=36 trials per training class, SD=2.44, minimum=29, maximum=39). We then identified the three feature numbers associated with each of the optimal training policies and averaged the decoding probabilities for those number labels on each test trial.

Using this approach, we examined whether average decoding evidence for the three features anticipated under each of the optimal training policies was higher than chance (8.33% based on the 12 possible feature numbers). Like the earlier section on policies, decoding evidence was averaged from +2.5s to +12.5s following test trial onset. Feature information associated with the more rewarding training policy was not detected on test tasks (OTC: M=8.71%, SD=1.12, *t*(37)=2.04, corrected *p*=0.392; MTL: M=8.37%, SD=0.40, *t*(37)=0.51, corrected *p*>0.99; OFC: M=8.28%, SD=0.52, *t*(37)=-0.60, corrected *p*>0.99; DLPFC: M=8.34%, SD=0.72, *t*(37)=0.05, corrected *p*>0.99). Equivalent results were found for features associated with the less rewarding training policy (OTC: M=8.61%, SD=0.91, *t*(37)=1.87, corrected *p*=0.486; MTL: M=8.38%, SD=0.59, *t*(37)=0.47, corrected *p*>0.99; OFC: M=8.37%, SD=0.52, *t*(37)=0.48, corrected *p*>0.99; DLPFC: M=8.38%, SD=0.57, *t*(37)=0.50, corrected *p*>0.99, Fig. S12).

## Discussion

This study aimed to investigate whether an SF&GPI algorithm could account for neural activity when humans transfer their experience from known to novel tasks. Behavioural results showed that human choices on new tasks relied on reusing policies that were successful in previous tasks. While this strategy was less optimal than a model-based process using a full model of the environment, past policies were applied in a reward-sensitive manner that led to high performance. These results were not due to a simple perseveration strategy (Fig. S5). An analysis of neural activity during test tasks showed that successful training policies were also represented in occipital-temporal and dorsolateral-prefrontal areas. These policies were prioritised as candidates for decision-making, with stronger activation than alternative policies that offered higher rewards. Activation strength in OTC was correlated with reuse behaviour. We found no evidence for the reactivation of features associated with successful training policies. Our results speak towards a role of OTC and DLPFC in implementing an efficient form of generalisation that is superior to model-free behaviour, but less optimal than full model-based computation operating on a complete model of the environment.

### Computational accounts of generalisation behaviour

Consistent with previous behavioural research (Tomov et al., 2021), a computational process based on SF&GPI could explain human generalisation performance. However, it did not capture participant choices perfectly. This was evident in data showing that participants made fewer choices leading to feature triplet ɸ(4) on test trials than the model predicted (Fig. 2G-H). Exploratory tests revealed that this was due to the presence of two distinct subgroups (Fig. S2). Half of the participants showed a full recapitulation of the SF&GPI predictions on test tasks. The other half showed a partial recapitulation. This suggests that some participants used different strategies on specific test tasks. When examining individual test tasks (Fig. S3), we further observed that the SF&GPI algorithm predicted the dominant choice in most cases. However, there was one test task on which choices were evenly split between the SF&GPI and MB predictions. Whereas most test tasks contained an anti-correlated structure in the feature weights that was similar to one or more training tasks, this outlier test task did not. This raises the possibility that at least some participants exploited structural similarities between the training and test tasks to determine their choices, similar to a Universal Value Function Approximator (UVFA) process in which similar task cues lead to similar rewards for a given action (Schaul et al., 2015). Tomov et al. (2021) observed evidence for UVFAs in a minority of subjects, but also had less structural overlap between training cues and the test cue. Future research should systematically manipulate the similarity between training and test scenarios to understand how this affects the computational process used for generalisation.

Post hoc behavioural predictions of a UVFA process are included in Figure S4. These predictions show consistencies with human choice, suggesting that UVFA could offer an alternative account of behavior. However, the neural findings from this experiment appear to be inconsistent with several UVFA predictions. Vanilla UVFAs predict equal activation of all available actions to compute and compare Q-values. In contrast, OTC showed activation of the optimal training actions/policies at choice onset, followed by sustained activation of the more rewarding action/policy (Fig. 3D), whereas DLPFC showed activation of the more rewarding action/policy (Fig. 3E). While this could be explained by a partial Markov decision process, in which people only evaluate actions that were reinforced during training, it would be inconsistent with the UVFA-like empirical choice profile on test task w=[1,1,-1] (Fig. S4B), in which a non-reinforced action is selected with high frequency. A UVFA explanation of the empirical choice profile on test task w=[1,1,-1] (Fig. S4B) would also predict approximately equal neural activation of both the objective best policy and the more rewarding training policy. Instead, we find that the more rewarding training policy is activated significantly more than the objective best policy, for which decoding evidence is at chance (Fig. S9B). Despite these neural inconsistencies, the behavioural similarities to UVFAs suggest that humans may utilise some kind of UVFA-like generalisation, perhaps in addition to SF&GPI-like policy reuse, in which the evaluated actions are not explicitly represented in the brain regions considered here. Exploring potential hybrid strategies that combine UVFAs with SF&GPI could be a promising area of future investigation.

Recent studies have also shown behavioural and neural evidence for reinforcement learning (RL) based on configurations of cues, over and above their individual elements (Duncan et al., 2018; Ballard et al., 2019). Applied to our setup, this would correspond to learning unique values for each (city, task) configuration. This account would not immediately capture the generalisation effects reported here. While an agent learning to respond to each configuration would be expected to learn optimal solutions to the training tasks, it would not have an obvious mechanism to map this knowledge to new tasks (i.e. new configurations of cues), the same issue facing a model free-agent (Fig. S5). Nonetheless, configural accounts could in principle be extended to support generalisation to novel configurations using function approximation. This would allow an agent to smoothly interpolate between the values for different (city, task) configurations experienced during training, to compute approximate values for the new (city, task) configurations on test trials. Interestingly, this is precisely what UVFAs do. While our experiment was designed to disambiguate between SF&GPI and MB control using a complete model, we do see signatures of UVFA-like response patterns (Fig. S4B), even though our neural results appear to be inconsistent with UVFA predictions (Fig. S9B). Exploring UVFAs as a generalisation of configural RL in the multi-task domain could be an interesting direction for future investigation.

A final account of generalisation behaviour could be a heuristic strategy, in which participants focus exclusively on the feature with the highest positive reward weight. Participants could then retrieve a policy from the training tasks that shared this top ranked feature. Such a heuristic would lead to SF&GPI-like behaviour in three test cases. However, it would not predict neural activation of the less rewarding training policy, as there would be no need to consider other features or policies once the top-ranked gem had been identified. Therefore, this heuristic strategy would not explain the neural effects in OTC, where both the more and less rewarding training policies were activated on test tasks (Fig. 3C-D).

### Neural predictions of SF&GPI

Consistent with SF&GPI predictions, we observed neural prioritisation of the optimal training policies during test tasks. This was most prominent in OTC. Evidence from our data suggests that two distinct sources contributed to decoding in this region. One was a reactivation of optimal training policies after seeing the test cue, as optimal training policies could be decoded earlier than what would be expected based on training trials. This could reflect expectations about upcoming stimuli in visual cortex (Aitken et al., 2020; Kok et al., 2014), with added modulation based on behavioural relevance. The other source was prioritised processing of the optimal training policies shown on the response screen. This enhanced processing could reflect value-driven attentional capture, in which stimuli previously associated with reward are preferentially attended to in new contexts (Anderson et al., 2011). While OTC is not a usual candidate for experiments on RL and decision-making, we note recent reports that neural replay decoded from OTC relates to the formation of successor representations (Wittkuhn et al., 2024), which are one of the theoretical foundations of SF&GPI. Neural prioritisation was also observed DLPFC, an area proposed to encode policies (Botvinick & An, 2008; Fine & Hayden, 2022). This finding aligns with DLPFC’s role in context-dependent behaviour (Badre & Nee, 2017; Flesch et al., 2022; Frith, 2000; Jackson et al., 2021; Rowe et al. 2000), as the task cue on each trial can be seen as a context cue that determines the current response mapping. One interpretation of our findings is that DLPFC can generalise this role outside a set of training cues, retrieving relevant response mappings for novel context cues with similar but non-identical structure to earlier contexts. We also note that although the prioritisation seen in our data is consistent with an SF&GPI-like process, it could also support hybrid models that use cached option values to identify useful candidates for particular decisions, and then perform model-based planning on that subset of options (Cushman & Morris, 2015; Morris et al., 2021).

Based on proposals that the MTL and OFC serve as predictive maps that encode information about future states (De Cothi & Barry, 2020; Geerts et al. 2020; Muhle-Karbe et al., 2023; Stachenfeld et al. 2017; Wimmer & Büchel, 2019), we predicted that features expected under the optimal training policies would be detected in these regions on test trials. We did not find evidence that this was the case. One possibility is that the features were not central to participants’ decision process, which could occur if choices were primarily based on structural similarities between the training and test cues. It is also possible that the features were used but that we were unable detect them. This could occur if feature numbers were represented with different neural patterns in the localiser task used to train the neural decoders than the main task. Cross-validated tests with policy decoders imply some support for this possibility, suggesting that decoders trained and tested on the main task tended to perform better, compared to those trained on the localiser and then tested on the main task (Fig. S13). Future experiments should therefore develop designs in which feature information can be remapped between the optimal and suboptimal policies within each block of a single paradigm to train more sensitive feature decoders. A second reason for not detecting features could be because each test task was repeated five times per block without feedback. This could have meant that feature triplets were used to compute expected rewards during initial test trials but that a policy was cached and used for remaining test trial repeats. Decoding predictive feature representations remains a critical target in future neural tests of SF&GPI.

It is also possible that OFC and MTL were engaged during the experiment but encoded other variables. For instance, one influential view of OFC function is that the OFC represents the prospect of reward for different options during decision-making (Kahnt et al. 2010, Knudsen & Wallis, 2022). The present experiment was not optimised to decode value signals due to the wide variation and largely unsystematic distribution of the rewards across training and test tasks (Fig. 4), a compromise that arose from the other factors we needed to balance in the experiment to test our central predictions. Some studies also suggest that hippocampal signalling may modulate the reinstatement strength of neural patterns in visual areas (Bosch et al., 2014; Hindy et al. 2016). Based on these factors, one speculative possibility could be that value information in OFC and hippocampal signalling modulate OTC and DLPFC activity on new tasks, in favour of successful past solutions.

**Figure 4.**
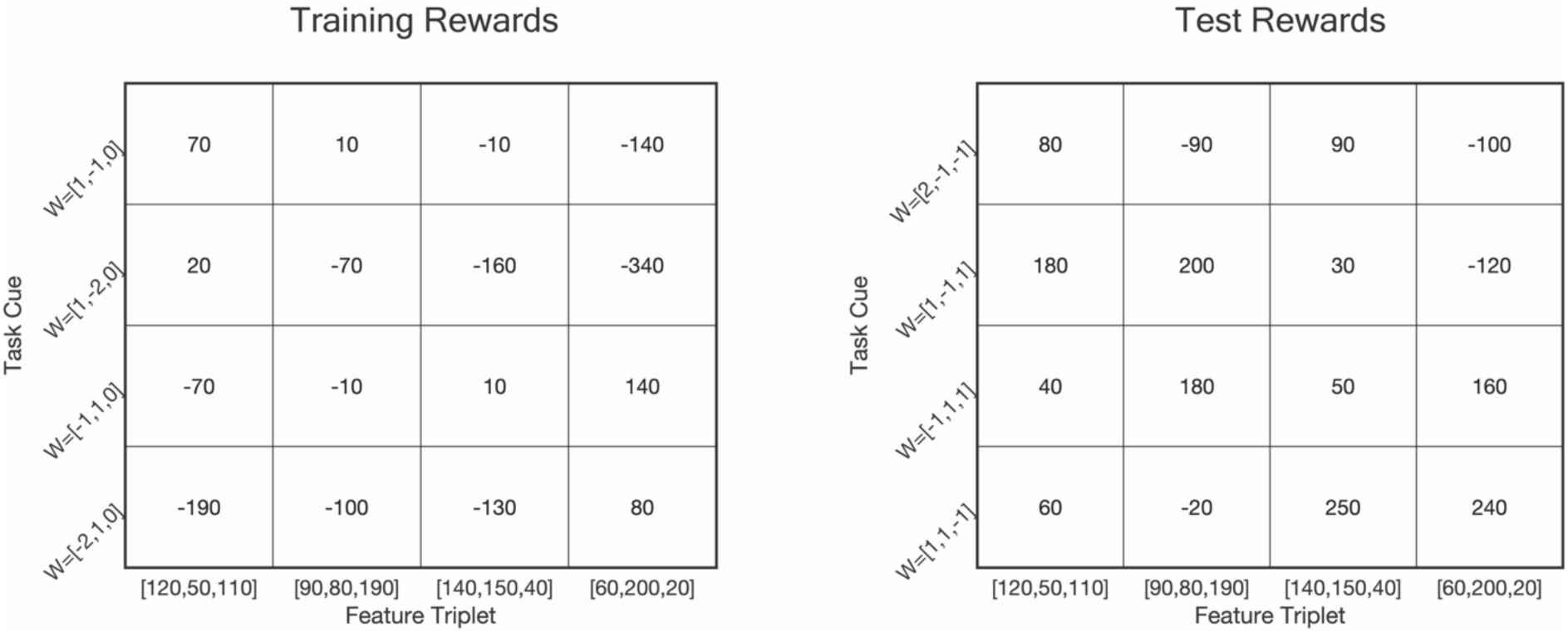
Reward outcomes for each combination of task cue and policy in the experiment. Rewards for training tasks are shown in the left-hand matrix. Rewards for test tasks are shown in the right-hand matrix. The order of feature triplets here is the same as other figures in the main text and supplement. [120,50,110] (or a permutation of these numbers) corresponds to ‘Feature Triplet 1’, [90,80,190] (or a permutation of these numbers) corresponds to ‘Feature Triplet 2’ and so on. Materials to reproduce this figure are available at https://gin.g-node.org/sam.hall-mcmaster/sfgpi-behavioural-analysis.

### Limitations

The present study had two main limitations. One was that each trial involved a single decision step. This meant that multi-step successor features were equivalent to single-step state features in the present design. This simplification was made to meet the practical constraints of fMRI scanning but future studies will need to devise practical ways to retain a multi-step element that can distinguish between computational processes using successor features and state features for generalisation. This would also help to distinguish reuse based on SF&GPI from more subtle forms of MB control that operate on a partial model of the environment. The most direct data we have on whether participants had a partial model in the current experiment were feature estimates collected after fMRI scanning. Specifically, participants were asked to estimate the features (gem numbers) for each city in the final experimental block (Fig. S6). If participants had a partial model of the environment, one prediction is that participants should have more accurate knowledge about features associated with the optimal training policies. We did not detect robust evidence for this prediction (corrected *p*-values>0.09, Fig. S6A). However, we did observe a significant correlation between having less accurate knowledge about features associated with the suboptimal training policies and the reuse of successful past solutions during test tasks (Spearman’s *Rho*=0.493, *p*=0.016, Fig. S6B). One possibility based on this result is that participants with higher policy reuse had a partial model of the environment or noisier memory for certain features. Unfortunately, the present design can only arbitrate between a computational process based on SF&GPI and a MB process that uses a complete model of the environment. It is unable to arbitrate between SF&GPI and forms of MB processing that use a partial model of the environment or have noisy memory. Multi-step designs could better address the separation between SF&GPI and partial MB control in future experiments. An MB agent with a partial model should be able to report features for each individual state that comprises the optimal training policies. A memory-based SF&GPI agent is expected to fail on such queries, as it stores a compact summary of this information that does not have individual state resolution. A second limitation was that the reward for the objective best policy was only 10-20 points higher than the more rewarding training policy (from among the optimal training policies). While we observed a high degree of reuse as predicted under an SF&GPI algorithm, it could be that the computational process would differ – with more equal evaluation across all options – if the reward prospect for previously unsuccessful policies had been higher at test. This would align with evidence that more computationally intensive forms of decision-making, such as MB planning, are adopted when the reward for accurate performance is heightened (Kool et al., 2017). The present experiment was not optimised to address meta-control between computational processes for different reward-effort trade-offs. Future research could therefore manipulate the difference in reward between successful and unsuccessful training policies during test tasks, to better understand the conditions under which an SF&GPI-like transfer process is (or is not) used.

## Conclusions

Overall, the present study provides behavioural and neural evidence that generalisation to new tasks was more consistent with an SF&GPI-based algorithm than an MB algorithm using a full model of the environment. These results do not rule out an MB algorithm operating on a partial model of the environment, or with noisy memory for features associated with suboptimal training choices. SF&GPI and UVFAs should also be systematically compared in the presence and absence of structural similarity between training and test cues in future studies. Ultimately, this experiment found that successful past solutions were prioritised as candidates for decision making on tasks outside the training distribution. This prioritisation provides flexibility when faced with new decisions problems and has lower computational cost than considering all available options. These findings take a step towards illuminating the flexible yet efficient nature of human intelligence.

## Methods

### Ethics Statement

The experimental protocol was approved by the ethics commission at the German Psychological Society (reference: SchuckNicolas2022-10-24AM) and conducted in accordance with the Declaration of Helsinki. Participants signed informed consent before each test session.

### Participants

Forty people participated in the experiment. One participant was excluded from the final sample due to low behavioural performance. The number of points earned in each scanning session was more than 2.5 standard deviations below the sample mean. Another participant was excluded due to excessive head motion. This was based on framewise displacement, which measures the change in head position between adjacent data points (Jenkinson et al., 2002). All functional runs in the second scanning session for the participant were more than 2.5 standard deviations above the sample mean for framewise displacement. The resulting 38 participants were between 18-35 years of age (mean=26 years, 23 female). All individuals had normal or corrected-to-normal vision and did not report an on-going neurological or psychiatric illness. €75 was paid for completing the experiment and €25 extra could be earned as a performance dependent bonus (€10 in session 1 and €15 in session two).

### Materials

Psychopy3 (RRID: SCR_006571, version 2021.2.3, Peirce et al., 2019) and Pavlovia (RRID:SCR_023320, https://pavlovia.org/) were used to prescreen prospective participants. Stimulus presentation during the scan sessions was controlled using Psychophysics Toolbox-3 (RRID:SCR_002881, version 3.0.17) in MATLAB (RRID:SCR_001622). The sessions used stimuli from Saryazdi et al. (2018) and city images from Rijan Hamidovic, Aleksander Pesaric, Pixabay and Burst that were retrieved from https://pexels.com. Scan preparation was completed on a computer outside the scanner with a spatial resolution of 2048 × 1152 and refresh rate of 60Hz (MATLAB version R2021b). The main experimental tasks were run on a stimulus PC with a spatial resolution of 1024 × 768 and a refresh rate of 60Hz, which projected to a magnetically safe screen for participants inside the scanner (MATLAB version R2017b). Responses were recorded using two fiber optic button boxes (Current Designs, Philadelphia, PA, USA). Participants used two buttons on the left box (middle finger and index finger) and three buttons on the right box (index finger, middle finger and ring finger). Output files from the scanner were converted to the BIDs naming convention (RRID:SCR_016124) using ReproIn (RRID:SCR_017184, HeuDiConv version 0.9.0).

Preprocessing and quality control were conducted using fMRIprep (Esteban et al., 2019, RRID:SCR_016216, version 20.2.4) and MRIQC (Esteban et al., 2017, RRID:SCR_022942, version 0.16.1). Behavioural analyses were performed in MATLAB (version 2021b). Neural analyses were performed in Python (RRID:SCR_008394, version 3.8.13), primarily using SciPy (RRID:SCR_008058, version 1.7.3), Pandas (RRID:SCR_018214, version 1.4.4), NumPy (RRID:SCR_008633, version 1.21.5), Matplotlib (RRID:SCR_008624, version 3.5.2), MNE (RRID:SCR_005972, version 1.4.2), Sklearn (RRID:SCR_019053, version 1.3.0) and Nilearn (RRID:SCR_001362, version 0.10.1). Data and code management was performed using DataLad (RRID:SCR_003931, version 0.17.9) and GIN (RRID:SCR_015864).

### Data and Code Access

Data and code for this project are openly available in the following repositories.

**Table 1.**
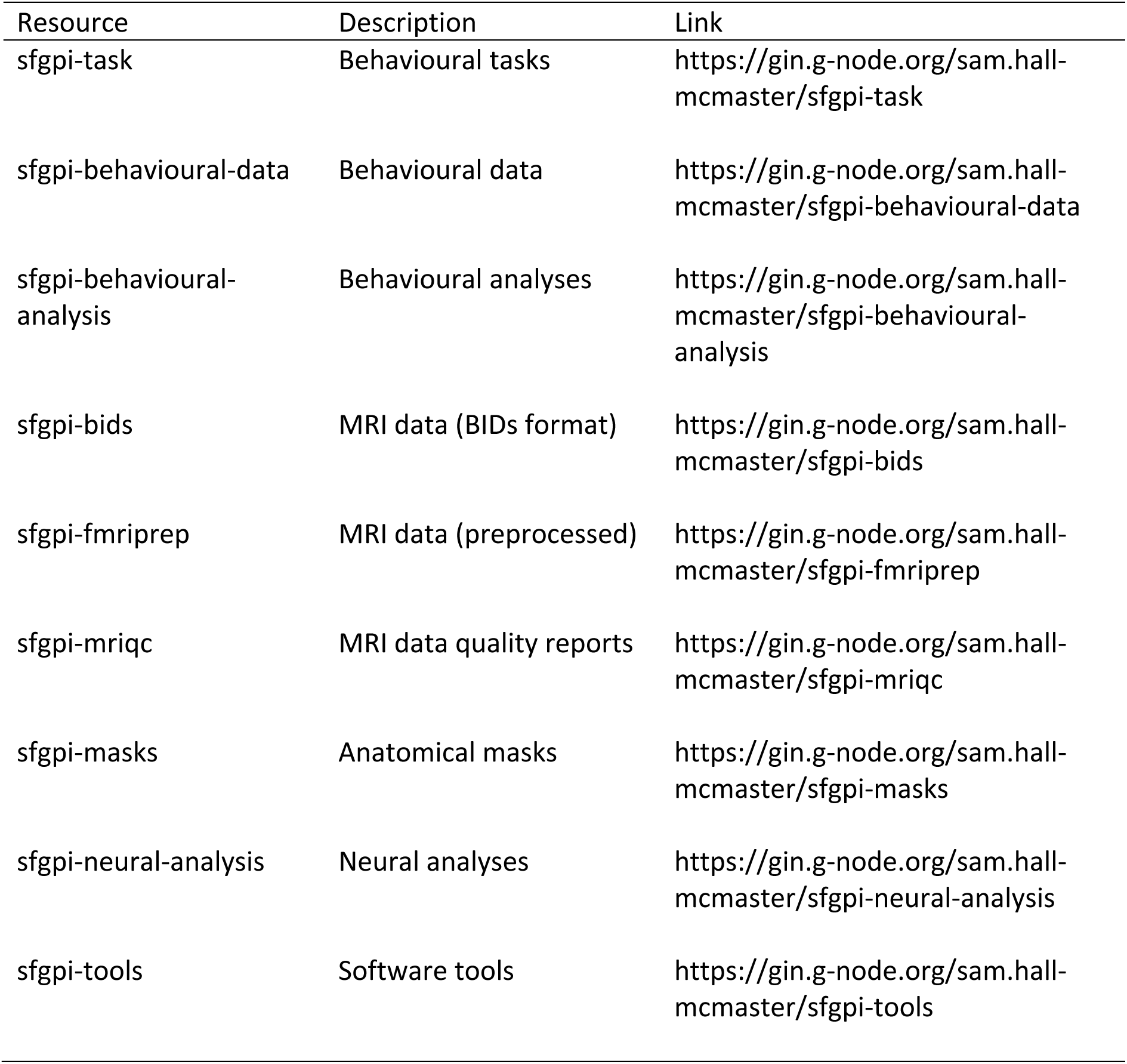
Data and code resources.

### Online Prescreening

Participants completed an online prescreening task prior to study admission. This was conceptually similar to the main task in MRI session two. However, it had different stimuli, different state features and a different theme to avoid biasing participants during scanning. The market values used for the prescreening were: w_train_ = {[1, −1, 0], [−1, 1, 0], [1, −2, 0], [−2, 1, 0]}. The state features were: ɸ = {[100, 0, 0], [30, 30, 160], [100, 100, 0], [0, 100, 70]}.

The vector elements in each triplet were shuffled with a randomly selected permutation rule for each participant. The theme used for prescreening was from Tomov et al. (2021). Participants completed 80 trials with equal numbers of each training task (w_train_) in a random order. No test tasks (w_test_) were presented. Trials had the same structure as Figure 1A but with different event timing. Market values were shown for 2.5s, no response limit was imposed, and feedback was shown for 3s. The cue-target interval was set to 0s and the inter-trial interval to 1s. Prospective participants needed to achieve an average reward above zero points to pass the prescreening and participate in the experiment. Up to three attempts could be made. 95% of people who passed the prescreening did so on their first attempt. Four prospective participants did not pass and were not admitted to the experiment.

### Session One

During the first experimental session, participants completed an associative retrieval task inside the MRI scanner. Session one was designed to produce a localiser dataset that could be as training data for neural decoders. Data from this session were used to train feature decoders but not policy decoders in the present paper. For the procedure behind the gem collector game, see session two.

#### Scan Preparation

In preparation for the first MRI scan, participants learned associations between cartoon objects (e.g. a balloon) and cities or number stimuli that would appear during session two. There were 4 cities (Sydney, Tokyo, New York, London) and 12 numbers (20, 40, 50, 60, 80, 90, 110, 120, 140, 150, 190, 200) in total. Each target was associated with two cartoon images, resulting in 32 associations. The associations between stimuli were randomly determined for each person. Participants completed a structured training procedure to learn the associations. The training began with the 8 object-city associations. On each trial, participants were shown a cartoon object (e.g. a balloon) for 1.5s, followed by a 0.25s blank delay. The four cities were then shown on screen in a random order. Participants needed to press the key (D, F, J, K, or L) corresponding to the screen position of the correct city within 5s. Following a response, feedback was shown for 2s (‘Correct!’ or ‘Incorrect!’). If the response was incorrect, participants were shown the correct object-city pairing. The trial concluded with a blank inter-trial interval (ITI) lasting 1s. Each block had 8 trials in total. This included one trial per association (e.g. four cities with two associated objects each), presented in a random order. Participants continued to complete blocks until achieving 8/8 correct responses. To reduce training time, the object duration on screen was shortened from 1.5s to 0.5s after the first block for each set of 8 associations. Using the same training structure, participants then learned 8 associations for each of the following number sets in turn: [20, 40, 50, 60], [80, 90, 110, 120], [140, 150, 190, 200]. To conclude the training, participants were tested on longer blocks with one trial for each of the 32 associations. Blocks continued until participants scored at least 90% correct (29/32 trials). After passing the 90% criterion once, participants had to do so one more time but with a shorter response deadline of 1.5s.

#### Scan Task

The MRI task for session one was a cued retrieval task. The setup was similar to the final training stage outlined in the previous section. On each trial, participants were shown a cartoon object for 0.5s and asked to imagine its associated target (i.e. the city or number associated with it). Following a blank delay, participants were presented with four possible response options in a random order on screen and needed to select the associated item within 1.5s. A feedback screen was then shown and the trial concluded with a blank ITI. The delay between the cartoon object offset and the response phase onset was drawn from a truncated exponential distribution, with a mean of 3s (lower bound=1.5s, upper bound=10s). Feedback (‘+10’ or ‘+0’) was shown for 0.5s. The ITI was drawn from a truncated exponential distribution with a mean of 2s (lower bound=1.5s, upper bound=10s). Participants completed 64 trials per block and 10 blocks in total. The trial order was constrained so that the correct target was not repeated on consecutive trials. Trials could also be labelled based on the following categories: cities (Sydney, Tokyo, New York, London), number set 1 (20, 40, 50, 60), number set 2 (80, 90, 110, 120) and number set 3 (140, 150, 190, 200). Consecutive trials with the same category were allowed to occur a maximum of two times per block. To motivate good performance, participants were shown their average accuracy, median RT and their bonus earnings up to that point at the end of each block. Fieldmaps and an T1w anatomical image were collected after the 5^th^ block. This scan resulted in 640 retrieval trials, with 40 trials for each target stimulus.

### Session Two

During the second experimental session, participants completed the gem collector game described in the main text. Sessions were held on consecutive days, with session two occurring the day after the session one.

#### Scan Preparation

Participants completed a brief preparation task to orient them to the gem collector theme and trial events. This included 30 practice trials. Practice trials had the same event structure as those shown in Figure 1. The market values used in the preparation task were: w_train_ = {[1, −1, 0], [−1, 1, 0]}. The state features were: ɸ = {[100, 0, 0], [30, 30, 160], [100, 100, 0], [0, 100, 70]}. The vector elements in each triplet were shuffled with a randomly selected permutation rule for each participant. The assignment between cities and state feature triplets was also randomised. Participants were told how the game profit was calculated on each trial and had to correctly calculate the profit for three example trials to confirm their understanding. The first 10 practice trials provided feedback after each choice. The remaining 20 trials consisted of 15 trials with feedback and 5 trials without feedback. Event timing was reduced across practice to prepare participants for the event speed during scanning. Market values were presented for 3.5s and a 5s response deadline was imposed for the first 10 trials. This was reduced to 2s and 1.5s for the remaining trials. The feedback (2.5s), cue-target interval (1s) and inter-trial interval (2s) had consistent durations. Participants did not need to reach a performance threshold on the practice trials to proceed to the scan task.

#### Scan Task

The cognitive task used in the second scanning session is described in the main text. On each trial, participants were shown the selling prices of three gem stimuli (2s). This was followed by a jittered blank delay. The specific duration on each trial was drawn from a truncated exponential distribution with a mean of 3.5s, a lower bound of 3s and an upper bound of 10s. Four city stimuli then appeared on screen in a random order. The random ordering prevented participants from preparing a motor response before seeing the choice stimuli. Participants had up 1.5s to choose a city. Upon entering a response, a feedback screen was presented (2.5s) that showed the selling prices, the number of gems in the selected city and the profit (or loss). The feedback screen was omitted on certain trials (described below). In the event that no response was made within 1.5s, ‘Too slow!’ was printed to the screen as feedback. The trial concluded with a jittered ITI. The specific duration on each trial was drawn from a truncated exponential distribution with a mean of 3s, a lower bound of 2.5s and an upper bound of 10s.

Each event in the trial sequence had a specific function. The selling prices were used to define the ‘task’ on each trial. A task is a context that specifies which features of the environment should be prioritised during decision making. For example, if square gems have a high selling price and the other gems sell for $0 per unit, the task on that trial is to choose the option with the highest number of square gems. In this way, the selling prices (which were numeric weight vectors) allowed us to manipulate the task on each trial in a precise manner. Another important trial event was the feedback. Each city was had unique number of gems. For example, Sydney might have 120 square gems, 50 triangle gems and 110 circle gems. The number of gems were state features (ɸ) associated with each city. The following set was used during scanning: ɸ = {[120, 50, 110], [90, 80, 190], [140, 150, 40], [60, 200, 20]}. The profit (or loss) on each trial was the dot product of the task weight triplet (the selling prices) and the state feature triplet (the gem numbers in the selected city).

The experiment had two main trial types. Training trials allowed participants to learn about the state features for each city because they showed feedback after each choice. Test trials were used to assess participants’ generalisation strategy and did not provide feedback. Participants were informed that the reward from all trials counted towards the €15 performance bonus in the session, even if feedback was not shown. Training and test trials were also distinguished based on the tasks (selling prices) shown to participants. The training tasks used in the experiment were: w_train_ = {[1, −1, 0], [−1, 1, 0], [1, −2, 0], [−2, 1, 0]}. The test tasks were: w_test_ = {[2, -1, -1], [-1, 1, 1], [1, -1, 1], [1, 1, -1]}. The reward for enacting each policy in each of the tasks is shown in Figure 4.

Each block had 68 trials. The order of training and test trials were pseudo-randomised based on a set of constraints. The block was divided into two phases. The first 32 trials were training trials (phase 1). The remaining 36 trials were a mixture of 16 training trials and 20 test trials (phase 2). Trials in the first phase were constrained so that: 1) equal numbers of the four training tasks were shown, 2) the same task was not repeated on consecutive trials and 3) trials with a particular optimal choice under SF&GPI (e.g. Sydney) were preceded by an equal number of trials with the same or a different optimal choice (e.g. 50% Sydney and 50% New York). Equivalent constraints were applied to the remaining trials (i.e. phase 2). Trials in phase 2 had the additional constraints that: 4) the phase did not begin with a test trial and 5) no more than 3 test trials were presented in a row. The two phases in each block were recorded as separate fMRI runs to aid leave-one-run-out cross-validation (described in the section on neural analyses).

Participants completed 6 blocks in total (12 fMRI runs). There were six possible permutation rules that could be used to order the numeric elements within each task and state feature triplet: {[1, 2, 3], [1, 3, 2], [2, 1, 3], [2, 3, 1], [3, 1, 2], [3, 2, 1]}. One configuration was used for each block. The configuration sequence across blocks was randomised. The state features for each city were also varied across blocks. We ensured that each city was paired with each state feature triplet at least once. A random mapping was used for the remaining two blocks. The mapping was pseudo-randomised across blocks so that no city had the same state feature triplet in consecutive blocks. The screen position of the three gem stimuli was randomised for each participant but remained consistent throughout the experiment. Permuting the triplet elements and changing the state features in each city across blocks preserved the logical structure of the experiment, while giving the appearance of new game rounds for participants.

#### Post Scan

To conclude session two, participants were asked to estimate the number of triangle, square and circle gems that had appeared in each city during the final block of the scan task. Estimates were restricted to a range between 0-250 gems. Participants were also asked to rate their confidence for each estimate on a scale from 0 (not confident at all) to 100 (completely confident). These measures were not probed during the scan itself to avoid biasing participants’ learning strategy.

### MRI Parameters

MRI acquisition parameters were based on Wittkuhn et al. (2021). Data were acquired using a 32-channel head coil on a 3-Tesla Siemens Magnetom TrioTim MRI scanner. Functional measurements were whole brain T2* weighted echo-planar images (EPI) with multi-band acceleration (voxel size=2×2×2xmm; TR=1250ms; echo time (TE)=26ms; flip angle (FA)=71 degrees; 64 slices; matrix = 96 × 96; FOV = 192 × 192mm; anterior-posterior phase encoding direction; distance factor=0%; multi-band acceleration=4). Slices were tilted +15 relative to the corpus callosum, with the aim of balancing signal quality from MTL and OFC (Weiskopf et al., 2006). Functional measurements began after five TRs (6.25s) to allow the scanner to reach equilibrium and help avoid partial saturation effects. Up to 401 volumes (session one) or 333 volumes (session two) were acquired during each task run; acquisition was ended earlier if participants had completed all trials. Fieldmaps were measured using the same scan parameters, except that two short runs of 20 volumes were collected with opposite phase encoding directions. Fieldmaps were later used for distortion correction in fMRIPrep (Esteban et al., 2019, RRID: SCR_016216, version 23.0.2). Anatomical measurements were acquired using T1 weighted Magnetization Prepared Rapid Gradient Echo (MPRAGE) sequences (voxel size=1×1×1xmm; TR=1900ms; TE=2.52ms; flip angle FA=9 degrees; inversion time (TI)=900ms; 256 slices; matrix = 192 × 256; FOV = 192 × 256mm).

### MRI Preprocessing

Results included in this manuscript come from preprocessing performed using *fMRIPrep* 23.0.2 (Esteban et al. (2019); Esteban et al. (2018); RRID:SCR_016216), which is based on *Nipype* 1.8.6 (K. Gorgolewski et al. (2011); K. J. Gorgolewski et al. (2018); RRID:SCR_002502).

#### Preprocessing of B_0_ Inhomogeneity Mappings

A total of 2 fieldmaps were found available within the input BIDS structure per subject. A *B_0_*-nonuniformity map (or *fieldmap*) was estimated based on two (or more) echo-planar imaging (EPI) references with topup (Andersson, Skare, and Ashburner (2003); FSL 6.0.5.1:57b01774).

#### Anatomical Data Preprocessing

A total of 2 T1-weighted (T1w) images were found within the input BIDS dataset per subject. All of them were corrected for intensity non-uniformity (INU) with N4BiasFieldCorrection (Tustison et al. 2010), distributed with ANTs 2.3.3 (Avants et al. 2008, RRID:SCR_004757). The T1w-reference was then skull-stripped with a *Nipype* implementation of the antsBrainExtraction.sh workflow (from ANTs), using OASIS30ANTs as target template. Brain tissue segmentation of cerebrospinal fluid (CSF), white-matter (WM) and gray-matter (GM) was performed on the brain-extracted T1w using fast (FSL 6.0.5.1:57b01774, RRID:SCR_002823, Zhang, Brady, and Smith 2001). An anatomical T1w-reference map was computed after registration of 2 T1w images (after INU-correction) using mri_robust_template (FreeSurfer 7.3.2, Reuter, Rosas, and Fischl 2010).

Brain surfaces were reconstructed using recon-all (FreeSurfer 7.3.2, RRID:SCR_001847, Dale, Fischl, and Sereno 1999), and the brain mask estimated previously was refined with a custom variation of the method to reconcile ANTs-derived and FreeSurfer-derived segmentations of the cortical gray-matter of Mindboggle (RRID:SCR_002438, Klein et al. 2017). Volume-based spatial normalization to two standard spaces (MNI152NLin6Asym, MNI152NLin2009cAsym) was performed through nonlinear registration with antsRegistration (ANTs 2.3.3), using brain-extracted versions of both T1w reference and the T1w template. The following templates were selected for spatial normalization and accessed with *TemplateFlow* (23.0.0, Ciric et al. 2022): *FSL’s MNI ICBM 152 non-linear 6th Generation Asymmetric Average Brain Stereotaxic Registration Model* (Evans et al. (2012), RRID:SCR_002823; TemplateFlow ID: MNI152NLin6Asym), *ICBM 152 Nonlinear Asymmetrical template version 2009c* (Fonov et al. (2009), RRID:SCR_008796; TemplateFlow ID: MNI152NLin2009cAsym).

#### Functional Data Preprocessing

For each of the 22 BOLD runs found per subject (across all tasks and sessions), the following preprocessing was performed. First, a reference volume and its skull-stripped version were generated using a custom methodology of *fMRIPrep*. Head-motion parameters with respect to the BOLD reference (transformation matrices, and six corresponding rotation and translation parameters) are estimated before any spatiotemporal filtering using mcflirt (FSL 6.0.5.1:57b01774, Jenkinson et al. 2002). The estimated *fieldmap* was then aligned with rigid-registration to the target EPI (echo-planar imaging) reference run. The field coefficients were mapped on to the reference EPI using the transform. BOLD runs were slice-time corrected to 0.588s (0.5 of slice acquisition range 0s-1.18s) using 3dTshift from AFNI (Cox and Hyde 1997, RRID:SCR_005927). The BOLD reference was then co-registered to the T1w reference using bbregister (FreeSurfer) which implements boundary-based registration (Greve and Fischl 2009). Co-registration was configured with six degrees of freedom. Several confounding time-series were calculated based on the *preprocessed BOLD*: framewise displacement (FD), DVARS and three region-wise global signals. FD was computed using two formulations following Power (absolute sum of relative motions, Power et al. (2014)) and Jenkinson (relative root mean square displacement between affines, Jenkinson et al. (2002)). FD and DVARS are calculated for each functional run, both using their implementations in *Nipype* (following the definitions by Power et al. 2014). The three global signals are extracted within the CSF, the WM, and the whole-brain masks.

Additionally, a set of physiological regressors were extracted to allow for component-based noise correction (*CompCor*, Behzadi et al. 2007). Principal components are estimated after high-pass filtering the *preprocessed BOLD* time-series (using a discrete cosine filter with 128s cut-off) for the two *CompCor* variants: temporal (tCompCor) and anatomical (aCompCor). tCompCor components are then calculated from the top 2% variable voxels within the brain mask. For aCompCor, three probabilistic masks (CSF, WM and combined CSF+WM) are generated in anatomical space. The implementation differs from that of Behzadi et al. in that instead of eroding the masks by 2 pixels on BOLD space, a mask of pixels that likely contain a volume fraction of GM is subtracted from the aCompCor masks. This mask is obtained by dilating a GM mask extracted from the FreeSurfer’s *aseg* segmentation, and it ensures components are not extracted from voxels containing a minimal fraction of GM. Finally, these masks are resampled into BOLD space and binarized by thresholding at 0.99 (as in the original implementation). Components are also calculated separately within the WM and CSF masks. For each CompCor decomposition, the *k* components with the largest singular values are retained, such that the retained components’ time series are sufficient to explain 50 percent of variance across the nuisance mask (CSF, WM, combined, or temporal). The remaining components are dropped from consideration. The head-motion estimates calculated in the correction step were also placed within the corresponding confounds file. The confound time series derived from head motion estimates and global signals were expanded with the inclusion of temporal derivatives and quadratic terms for each (Satterthwaite et al. 2013). Frames that exceeded a threshold of 0.5 mm FD or 1.5 standardized DVARS were annotated as motion outliers.

Additional nuisance timeseries are calculated by means of principal components analysis of the signal found within a thin band (*crown*) of voxels around the edge of the brain, as proposed by (Patriat, Reynolds, and Birn 2017). The BOLD time-series were resampled into standard space, generating a *preprocessed BOLD run in MNI152NLin6Asym space*. First, a reference volume and its skull-stripped version were generated using a custom methodology of *fMRIPrep*. The BOLD time-series were resampled onto the following surfaces (FreeSurfer reconstruction nomenclature): *fsnative*. All resamplings can be performed with *a single interpolation step* by composing all the pertinent transformations (i.e. head-motion transform matrices, susceptibility distortion correction when available, and co-registrations to anatomical and output spaces). Gridded (volumetric) resamplings were performed using antsApplyTransforms (ANTs), configured with Lanczos interpolation to minimize the smoothing effects of other kernels (Lanczos 1964). Non-gridded (surface) resamplings were performed using mri_vol2surf (FreeSurfer).

Many internal operations of *fMRIPrep* use *Nilearn* 0.9.1 (Abraham et al. 2014, RRID:SCR_001362), mostly within the functional processing workflow. For more details of the pipeline, see the section corresponding to workflows in *fMRIPrep*’s documentation.

### Computational Models

Theoretical predictions for choices on test tasks were generated using code from Tomov et al. (2021). The code was adapted to the current task design, which included changing the training tasks, test tasks, state features and the state space. Model training procedures remained the same.

The models were situated in a standard reinforcement learning formalism called a Markov Decision Process *M* = (*S*, *A*, *p*, *r*, *γ*), where *S* is a set of states, *A* is a set of actions, *p*(*s*′| *s*, *a*) is the probability of transitioning to a subsequent state (s’) from an initial state (s) after taking action a, *r*(*s*) is the reward received when entering state *s*, and *γ* is a discount factor between 0 and 1 that down-weights future rewards. The environment also contained state features ɸ(s) and tasks *w*. Policies defining the action an agent takes in each state are denoted *π*.

#### Model Based (MB)

The MB algorithm had perfect knowledge about the environment. To decide what to do on each test task, it computed expected values for all possible actions. Each expected value was defined as:

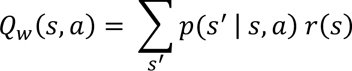

The experimental design used deterministic transitions and thus each entry in *p*(*s*′ | *s*, *a*) was 0 or 1. The reward, *r*(*s*), was computed as:

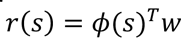

The MB algorithm selected the policy on the current test task with the highest expected value:

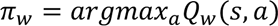

#### Successor Features and Generalised Policy Improvement (SF&GPI)

The SF&GPI algorithm is designed to learn estimates of the state features called successor features:

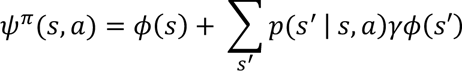

Since no further action could be taken in the terminal state (s’), this simplified to:

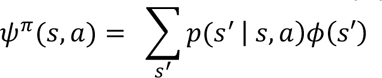

This simplification meant that successor features were equivalent to state features in the present design. The SF&GPI algorithm then performs generalised policy improvement, which computes a policy for the current test task based on earlier training policies. The first step in this process is to compute expected values on the current test task under a set of earlier policies {*π*_1_ … *π_n_*}:

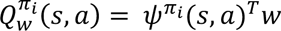

An interesting property of the SF&GPI algorithm is that it is flexible with respect to the policies it considers from training. When considering all policies experienced during training, an SF&GPI algorithm makes the same predictions as an MB algorithm in the present design. However, when generalised policy improvement is restricted to the optimal training policies, its predictions on test tasks differ:

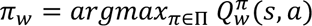

Here the set of policies, Π, contains the optimal training policies but not all policies. We used this formulation of SF&GPI to distinguish between a model of generalisation that evaluates optimal training solutions on test tasks and an MB generalisation process that evaluates all possible solutions.

### Behavioural Analyses

17 statistical tests are reported in the main text for behavioural data. The corresponding *p*-values reported were therefore corrected for 17 tests using the Bonferroni-Holm correction (Holm, 1979). Auxiliary tests presented in the supplement but mentioned in the main text were separately corrected using Bonferroni-Holm. Supplementary figure legends provide more details about the correction applied in these cases.

#### Average Performance

The average points per choice on training trials was computed for each participant. To compare this performance against a sensible baseline, we simulated an equivalent sample of 38 agents that made random choices on the same trials. To ensure comparability, points from random choices were replaced with 0 points for trials where the equivalent human participant omitted a response. The average points per choice was calculated for each agent. The two distributions (human and agent) were then compared using an independent two-tailed t-test. This process was repeated to assess average performance on the test trials.

#### State Analysis

The present experiment contained four terminal states. Each terminal state was defined by a unique feature triplet from the following set: ɸ = [120, 50, 110], [90, 80, 190], [140, 150, 40], [60, 200, 20]. The order of individual elements within each triplet varied across blocks in the experiment. The most rewarding terminal states during training trials were associated with ɸ(1) or ɸ(4). To assess whether participants made more choices leading ɸ(1) or ɸ(4), we recorded the proportion of choices leading to each terminal state across all training trials in the experiment. We then compared the proportions between each pair of states using paired two-tailed t-tests. The one exception was that we did not compare the choice numbers between ɸ(2) and ɸ(3), as this comparison was less related to the computational modelling predictions. To assess whether the same choice profile was recapitulated at test, the process above was repeated for test trials.

#### Reuse Behaviour

To further quantify the reuse of policies leading to ɸ(1) or ɸ(4) on test tasks, we computed the proportion of test trials in which choices to ɸ(1) or ɸ(4) were selected for each participant. To assess whether the proportions were above chance, the distribution was compared to a mean of 0.5 in a one sample two-tailed t-test. To understand whether policy reuse was sensitive to prospective rewards, we computed the proportion of test trials in which participants selected the more rewarding optimal policy from training. While the previous analysis counted choices to either ɸ(1) or ɸ(4) regardless of outcome, this analysis counted choices to ɸ(1) or ɸ(4) only when the more rewarding of the two was selected.

Since participants had a ¼ chance of selecting the more rewarding optimal training policy at random from the response array, the empirical proportions were compared to a mean of 0.25 in a one sample two-tailed t-test. The same process was used to quantify how often the optimal policy was selected on training trials.

### Neural Analyses

Decoding was performed separately for each ROI. The voxel activity on each trial was extracted from the current ROI and smoothed 2-4mm. The specific smoothing was determined in validation analyses that did not use the trials needed to test our hypothesis. The resulting data were detrended, z-scored and motion confounds (identified by fMRIPrep) were removed. Data cleaning was performed separately for each functional run using Nilearn signal clean. Trials without a response were then removed. The decoding procedure next entered a cross-validation loop. One block was held out to use as test data. The remaining blocks were used to train the decoder. A random sub-sample was taken to match the number of trials for each training class. It is important to note that only trials with feedback were used for decoder training (see Fig. 1A). Voxel activity 4-6s following feedback onset was extracted and the city shown on the feedback screen was used as the training label. The specific time lag was determined in validation analyses (see Fig. S7). The resulting voxel patterns and labels were used to train a logistic regression model. This was implemented in Sklearn with an L2 penalty term and a one-vs-the-rest multiclass strategy.

For more details, please see the Sklearn documentation. The trained decoder was then tested on each trial in the held-out run. Testing was performed at multiple TRs on each trial, from cue onset (TR_0_) through to 15s after it (TR_+12_). This resulted in one probability score per city stimulus at each TR in the held-out test trial. Each score was the probability – according to the model - that a particular city was the correct label for the neural data provided. The class probabilities at a specific data point did not need to sum to 1 (due to the one-vs-the-rest approach). The cross-validation procedure was repeated until all functional runs had been used as held-out test data. For robustness, the analysis procedure was repeated 100 times with random sub-sampling on each iteration. The results were based on model probabilities averaged across iterations. These probabilities are interpreted as decoding evidence throughout the paper.

#### ROIs

To extract voxel patterns from specific regions, we generated masks for the four ROIs using FreeSurfer cortical parcellations. Each mask was composed of one or more bilateral parcellations. Masks were specific to each participant. DLPFC was based on the rostral middle frontal gyrus parcellation. MTL combined the hippocampus, parahippocampal gyrus and entorhinal cortex parcellations. OFC combined the medial and lateral OFC parcellations. OTC combined the cuneus, lateral occipital, pericalcarine, superioparietal, lingual, inferioparietal, fusiform, inferiotemporal, parahippocampal and middle temporal parcellations. For more information, please see Desikan et al. (2006) and the FreeSurfer documentation: https://surfer.nmr.mgh.harvard.edu/fswiki/FsTutorial/AnatomicalROI/FreeSurferColorLUT

#### Validation

To validate the decoding pipeline, we trained neural decoders to classify the city observed during feedback on training trials. We then tested how well we could recover the chosen city on held out training trials. One functional run was held out at a time to assess decoding performance. As training trials were not related to our hypothesis, we used the validation procedure to explore three possible time shifts (+4s, +5s, +6s seconds following feedback onset) and three possible smoothing levels (0mm, 2mm, 4mm) that could be used to optimise decoder training. The validation results were based on 25 iterations. Data were subsampled at random on each iteration to match the number of trials from each class. The highest performing decoding parameters were selected for each ROI based on the validation results (Fig. S7). These parameters were later used when testing our hypothesis about neural reactivation during test tasks.

#### Policy Decoding

The choice on each test trial could fall into one of four categories: 1) the more rewarding policy (among the optimal training policies); 2) the less rewarding policy (among the optimal training policies); 3) the objective best policy; and 4) the remaining policy. To examine the neural evidence for these categories, we extracted decoding probabilities for the four choice stimuli on test trials – that is, on trials that did not show feedback (Fig. 1B). This extraction was first performed for the time point closest to cue onset (TR_0_). The four decoding probabilities on each test trial at this time point – one for each choice stimulus - were divided into the four categories above. To avoid contamination from the previous trial, probabilities were removed in cases where the policy category on the current test trial was the same as the previous trial. Once this operation had been performed for all test trials, the probabilities in each category were averaged over trials. This process was repeated at each time point until 15s after cue onset (TR_+12_). The output of this process was one decoding time course per policy category, per participant. These were stored in a 38 × 4 × 12 matrix, in which dimensions were participants x categories x time points (TRs). A separate matrix was generated for each ROI. To test the average decoding strength on test trials against chance, we averaged the matrices above from 2.5s after cue onset (TR_+2_) through to 12.5s (TR_+10_).

This time window was selected based on average test trial duration of 10s (Fig. 1B) and the hemodynamic delay. The 2.5s starting point was selected based on validation analysis (Fig. S7), which showed that an event often took two TRs to have a discernible impact on the decoding probabilities. The average probabilities for the optimal training policies were compared to the chance rate of 0.25, using one-sample two-tailed t-tests. The resulting p-values were corrected for 8 comparisons (2 policy categories x 4 ROIs) using the Bonferroni-Holm correction. To assess the evidence for the optimal training policies within OTC and DLPFC in a time resolved manner, we conducted one-sample cluster-based permutation tests against a population mean of 0.25 (window tested=2.5-12.5s) using mne.stats.permutation_cluster_1samp_test. To assess neural prioritisation in each ROI, we took the average decoding evidence for each of the optimal training policies and subtracted the average evidence for the objective best policy on test tasks. Prioritisation scores were tested against for significance using one-sample two-tailed t-tests. The two prioritisation scores within each ROI were further compared with two-sample two-tailed t-tests. The resulting p-values were corrected for 12 comparisons (2 prioritisation scores x 4 ROIs and 4 follow up tests) using the Bonferroni-Holm correction.

#### Control Analyses

We performed two control analyses to examine the policy results from OTC and DLPFC. To distinguish between neural reactivation and processing of options during the response screen, we re-ran the decoding analysis but tested the decoder on time points locked to the response phase onset. To be cautious about interpreting evidence at the onset itself, we set TR_0_ to be the earlier TR closest to each response phase onset (rather than rounding to the closest TR which could potentially contain signal further into the response phase). We then performed cluster-based permutation tests from 0s to +7.5s relative to the response phase onset. Trials with feedback were used to establish a baseline time lag for the decoding of policy information. This was contrasted with the time course observed on trials without feedback. To control for the impact of choice on the core results, we examined decoding evidence for the optimal training policies on test trials where participants did not select the corresponding policy. For example, when extracting decoding probabilities for the more rewarding policy (among the optimal training policies), we extracted probabilities from trials where participants selected a different policy. We then repeated statistical tests for the average decoding evidence and cluster-based permutation tests across time. The Bonferroni-Holm correction was applied for the four statistical tests comparing average decoding strength against chance (2 policy categories x 2 ROIs).

#### Feature Decoding

To decode feature information, logistic decoders were trained on fMRI data described in *Session One*. Trials in which participants selected the correct number target were eligible as training data. The response phase onset was taken for each eligible trial and time shifted 4-6s. The closest TR to this time point was used the neural training data from that trial. The training label was the selected number target. Training data was locked to the response phase (rather than feedback events for policy decoding) because feedback was only presented on error trials in session one. The time shift and spatial smoothing used were the same as policy decoding. After training, the logistic decoders were evaluated on each TR from the test trials from *Session Two.* This resulted in a separate decoding probability for the 12 feature numbers at each TR on each test trial. We then iterated over the test trials and extracted the decoding probabilities for the features associated with more (or less) rewarding training policy and averaged them. Feature probabilities were excluded from the average in cases where a feature number was the same as one of the optimal rewards from training. Probabilities were also excluded in cases where the features being decoded were the same as those selected on the previous trial. These control measures were intended to get an estimate of feature evidence that was independent from the previous trial and could not be explained as a reward prediction. The remaining procedure was the same as policy decoding. The extracted probabilities were averaged over test trials for each ROI and then averaged from 2.5s after cue onset through to 12.5s. The average probabilities for features associated with the optimal training policies were compared to a chance rate of 1/12, using one-sample two-tailed t-tests. The resulting p-values were corrected for 8 comparisons (2 policy categories x 4 ROIs) using the Bonferroni-Holm correction.

#### Associations with Reuse Behaviour

Spearman correlations were used to test whether policy decoding effects were associated with choices during the experiment. The average decoding evidence for the more rewarding training policy was correlated with the proportion of test trials on which participants made the choice predicted by SF&GPI. The decoding evidence had a precautionary measure to avoid contamination from the previous trial (see the earlier section on *Policy Decoding*). For consistency, the behavioural proportion used in correlation tests was calculated with the same trials used for the decoding evidence. The correlation was run twice, once for OTC and once for DLPFC. The process was then repeated but using neural prioritisation as the neural measure (the difference in decoding evidence for the more rewarding training policy and the objective best policy). The original process was then repeated one more time but using the proportion of model-based choices on test tasks as the behavioural measure. As correlation tests within each ROI did not use fully independent data, *p*-values were corrected for the two ROIs tested using the Bonferroni-Holm correction.

## References

Abraham, A., Pedregosa, F., Eickenberg, M., Gervais, P., Mueller, A., Kossaifi, J., … & Varoquaux, G. (2014). Machine learning for neuroimaging with scikit-learn. Frontiers in Neuroinformatics, 8, 14. 10.3389/fninf.2014.00014

Anderson, B. A., Laurent, P. A., & Yantis, S. (2011). Value-driven attentional capture. Proceedings of the National Academy of Sciences, 108(25), 10367–10371. 10.1073/pnas.110404710

Andersson, J. L., Skare, S., & Ashburner, J. (2003). How to correct susceptibility distortions in spin-echo echo-planar images: application to diffusion tensor imaging. Neuroimage, 20(2), 870–888. 10.1016/S1053-8119(03)00336-7

Aitken, F., Menelaou, G., Warrington, O., Koolschijn, R. S., Corbin, N., Callaghan, M. F., & Kok, P. (2020). Prior expectations evoke stimulus-specific activity in the deep layers of the primary visual cortex. PLoS Biology, 18(12), e3001023.

Avants, B. B., Epstein, C. L., Grossman, M., & Gee, J. C. (2008). Symmetric diffeomorphic image registration with cross-correlation: Evaluating automated labeling of elderly and neurodegenerative brain. Medical Image Analysis, 12(1), 26–41. 10.1016/j.media.2007.06.004

Badre, D., & Nee, D. E. (2018). Frontal cortex and the hierarchical control of behavior. Trends in Cognitive Sciences, 22(2), 170–188. 10.1016/j.tics.2017.11.005

Ballard, I. C., Wagner, A. D., & McClure, S. M. (2019). Hippocampal pattern separation supports reinforcement learning. Nature Communications, 10(1), 1073.

Barreto, A., Dabney, W., Munos, R., Hunt, J. J., Schaul, T., van Hasselt, H. P., & Silver, D. (2017). Successor features for transfer in reinforcement learning. Advances in neural information processing systems, 30.

Barreto, A., Borsa, D., Quan, J., Schaul, T., Silver, D., Hessel, M., … & Munos, R. (2018, July). Transfer in deep reinforcement learning using successor features and generalised policy improvement. In International Conference on Machine Learning (pp. 501–510). PMLR.

Barreto, A., Hou, S., Borsa, D., Silver, D., & Precup, D. (2020). Fast reinforcement learning with generalized policy updates. Proceedings of the National Academy of Sciences, 117(48), 30079–30087. 10.1073/pnas.1907370117

Behrens, T. E., Muller, T. H., Whittington, J. C., Mark, S., Baram, A. B., Stachenfeld, K. L., & Kurth-Nelson, Z. (2018). What is a cognitive map? Organizing knowledge for flexible behavior. Neuron, 100(2), 490–509. 10.1016/j.neuron.2018.10.002

Behzadi, Y., Restom, K., Liau, J., & Liu, T. T. (2007). A component based noise correction method (CompCor) for BOLD and perfusion based fMRI. Neuroimage, 37(1), 90–101. 10.1016/j.neuroimage.2007.04.042

Bosch, S. E., Jehee, J. F., Fernández, G., & Doeller, C. F. (2014). Reinstatement of associative memories in early visual cortex is signaled by the hippocampus. Journal of Neuroscience, 34(22), 7493–7500. 10.1523/JNEUROSCI.0805-14.2014

Botvinick, M., & An, J. (2008). Goal-directed decision making in prefrontal cortex: A computational framework. Advances in Neural Information Processing Systems, 21.

Carvalho, W., Tomov, M.S., de Cothi, W., Barry, C., & Gershman, S.J. (2024). Predictive representations: Building blocks of intelligence. Neural Computation (accepted)

Ciric, R., Thompson, W. H., Lorenz, R., Goncalves, M., MacNicol, E. E., Markiewicz, C. J., … & Esteban, O. (2022). TemplateFlow: FAIR-sharing of multi-scale, multi-species brain models. Nature Methods, 19(12), 1568–1571. 10.1038/s41592-022-01681-2

Cox, R. W., & Hyde, J. S. (1997). Software tools for analysis and visualization of fMRI data. NMR in Biomedicine, 10(4-5), 171–178. https://doi.org/10.1002/(SICI)1099-1492(199706/08)10:4/5<171::AID-NBM453>3.0.CO;2-L

Cushman, F., & Morris, A. (2015). Habitual control of goal selection in humans. Proceedings of the National Academy of Sciences, 112(45), 13817–13822. 10.1073/pnas.1506367112

Dale, A. M., Fischl, B., & Sereno, M. I. (1999). Cortical surface-based analysis: I. Segmentation and surface reconstruction. Neuroimage, 9(2), 179–194. 10.1006/nimg.1998.0395

De Cothi, W., & Barry, C. (2020). Neurobiological successor features for spatial navigation. Hippocampus, 30(12), 1347–1355. 10.1002/hipo.23246

Dekker, R. B., Otto, F., & Summerfield, C. (2022). Curriculum learning for human compositional generalization. Proceedings of the National Academy of Sciences, 119(41), e2205582119. 10.1073/pnas.2205582119

Desikan, R. S., Ségonne, F., Fischl, B., Quinn, B. T., Dickerson, B. C., Blacker, D., … & Killiany, R. J. (2006). An automated labeling system for subdividing the human cerebral cortex on MRI scans into gyral based regions of interest. Neuroimage, 31(3), 968–980. 10.1016/j.neuroimage.2006.01.021

Duncan, K., Doll, B. B., Daw, N. D., & Shohamy, D. (2018). More than the sum of its parts: a role for the hippocampus in configural reinforcement learning. Neuron, 98(3), 645–657.

Esteban, O., Markiewicz, C. J., Blair, R. W., Moodie, C. A., Isik, A. I., Erramuzpe, A., … & Gorgolewski, K. J. (2019). fMRIPrep: a robust preprocessing pipeline for functional MRI. Nature Methods, 16(1), 111–116. 10.1038/s41592-018-0235-4

Esteban, O., Markiewicz, C. J., Goncalves, M., Provins, C., Salo, T., Kent, J. D., DuPre, E., Ciric, R., Pinsard, B., Blair, R. W., Poldrack, R. A., & Gorgolewski, K. J. (2024). fMRIPrep: A robust preprocessing pipeline for functional MRI (23.2.1). Zenodo. 10.5281/zenodo.10790684

Esteban, O., Birman, D., Schaer, M., Koyejo, O. O., Poldrack, R. A., & Gorgolewski, K. J. (2017). MRIQC: Advancing the automatic prediction of image quality in MRI from unseen sites. PloS one, 12(9), e0184661. 10.1371/journal.pone.0184661

Evans, A. C., Janke, A. L., Collins, D. L., & Baillet, S. (2012). Brain templates and atlases. Neuroimage, 62(2), 911–922. 10.1016/j.neuroimage.2012.01.024

Fine, J. M., & Hayden, B. Y. (2022). The whole prefrontal cortex is premotor cortex. Philosophical Transactions of the Royal Society B, 377(1844), 20200524. 10.1098/rstb.2020.0524

Fischl, B. (2012). FreeSurfer. Neuroimage, 62(2), 774–781. 10.1016/j.neuroimage.2012.01.021

Flesch, T., Juechems, K., Dumbalska, T., Saxe, A., & Summerfield, C. (2022). Orthogonal representations for robust context-dependent task performance in brains and neural networks. Neuron, 110(7), 1258–1270. 10.1016/j.neuron.2022.01.005

Fonov, V. S., Evans, A. C., McKinstry, R. C., Almli, C. R., & Collins, D. L. (2009). Unbiased nonlinear average age-appropriate brain templates from birth to adulthood. NeuroImage, 47, S102. 10.1016/S1053-8119(09)70884-5

Frith, C. D. (2000). The role of dorsolateral prefrontal cortex in the selection of action as revealed by functional imaging. In S. Monsell & J. Driver (Eds.), Control of Cognitive Processes (pp. 544–565). MIT Press.

Geerts, J. P., Chersi, F., Stachenfeld, K. L., & Burgess, N. (2020). A general model of hippocampal and dorsal striatal learning and decision making. Proceedings of the National Academy of Sciences, 117(49), 31427–31437. 10.1073/pnas.2007981117

Gorgolewski, K., Burns, C. D., Madison, C., Clark, D., Halchenko, Y. O., Waskom, M. L., & Ghosh, S. S. (2011). Nipype: a flexible, lightweight and extensible neuroimaging data processing framework in python. Frontiers in Neuroinformatics, 5, 13. 10.3389/fninf.2011.00013

Gorgolewski, Krzysztof J., Oscar Esteban, Christopher J. Markiewicz, Erik Ziegler, David Gage Ellis, Michael Philipp Notter, Dorota Jarecka, et al. 2018. Nipype. Zenodo. 10.5281/zenodo.596855

Greve, D. N., & Fischl, B. (2009). Accurate and robust brain image alignment using boundary-based registration. Neuroimage, 48(1), 63–72. 10.1016/j.neuroimage.2009.06.060.

Haxby, J. V., Gobbini, M. I., Furey, M. L., Ishai, A., Schouten, J. L., & Pietrini, P. (2001). Distributed and overlapping representations of faces and objects in ventral temporal cortex. Science, 293(5539), 2425–2430. 10.1126/science.1063736

Hindy, N. C., Ng, F. Y., & Turk-Browne, N. B. (2016). Linking pattern completion in the hippocampus to predictive coding in visual cortex. Nature Neuroscience, 19(5), 665–667. 10.1038/nn.4284

Holm, S. (1979). A simple sequentially rejective multiple test procedure. Scandinavian Journal of Statistics, 65-70.

Jackson, J. B., Feredoes, E., Rich, A. N., Lindner, M., & Woolgar, A. (2021). Concurrent neuroimaging and neurostimulation reveals a causal role for dlPFC in coding of task-relevant information. Communications Biology, 4(1), 588. 10.1038/s42003-021-02109-x

Jenkinson, M., Bannister, P., Brady, M., & Smith, S. (2002). Improved optimization for the robust and accurate linear registration and motion correction of brain images. Neuroimage, 17(2), 825–841. 10.1006/nimg.2002.1132.

Kahnt, T., Heinzle, J., Park, S. Q., & Haynes, J. D. (2010). The neural code of reward anticipation in human orbitofrontal cortex. Proceedings of the National Academy of Sciences, 107(13), 6010–6015. 10.1073/pnas.0912838107

Klein, A., Ghosh, S. S., Bao, F. S., Giard, J., Häme, Y., Stavsky, E., … & Keshavan, A. (2017). Mindboggling morphometry of human brains. PLoS Computational Biology, 13(2), e1005350. 10.1371/journal.pcbi.1005350.

Knudsen, E. B., & Wallis, J. D. (2022). Taking stock of value in the orbitofrontal cortex. Nature Reviews Neuroscience, 23(7), 428–438. 10.1038/s41583-022-00589-2

Kok, P., Failing, M. F., & de Lange, F. P. (2014). Prior expectations evoke stimulus templates in the primary visual cortex. Journal of Cognitive Neuroscience, 26(7), 1546–1554.

Lanczos, C. (1964). Evaluation of noisy data. Journal of the Society for Industrial and Applied Mathematics, Series B: Numerical Analysis, 1(1), 76–85. 10.1137/0701007.

Luettgau, L., Erdmann, T., Veselic, S., Stachenfeld, K. L., Moran, R., Kurth-Nelson, Z., & Dolan, R. J. (2023, September 1). Decomposing dynamical subprocesses for compositional generalization. 10.31234/osf.io/sxn4a

Maris, E., & Oostenveld, R. (2007). Nonparametric statistical testing of EEG-and MEG-data. Journal of Neuroscience Methods, 164(1), 177–190. 10.1016/j.jneumeth.2007.03.024

Morris, A., Phillips, J., Huang, K., & Cushman, F. (2021). Generating options and choosing between them depend on distinct forms of value representation. Psychological Science, 32(11), 1731–1746. 10.1177/09567976211005702

Muhle-Karbe, P. S., Sheahan, H., Pezzulo, G., Spiers, H. J., Chien, S., Schuck, N. W., & Summerfield, C. (2023). Goal-seeking compresses neural codes for space in the human hippocampus and orbitofrontal cortex. Neuron, 111(23), 3885–3899. 10.1016/j.neuron.2023.08.021

Patriat, R., Reynolds, R. C., & Birn, R. M. (2017). An improved model of motion-related signal changes in fMRI. Neuroimage, 144, 74–82. 10.1016/j.neuroimage.2016.08.051

Peirce, J. W., Gray, J. R., Simpson, S., MacAskill, M. R., Höchenberger, R., Sogo, H., Kastman, E., Lindeløv, J. (2019). PsychoPy2: Experiments in behavior made easy. Behavior Research Methods. 10.3758/s13428-018-01193-y

Power, J. D., Mitra, A., Laumann, T. O., Snyder, A. Z., Schlaggar, B. L., & Petersen, S. E. (2014). Methods to detect, characterize, and remove motion artifact in resting state fMRI. Neuroimage, 84, 320–341. 10.1016/j.neuroimage.2013.08.048

Rowe, J. B., Toni, I., Josephs, O., Frackowiak, R. S., & Passingham, R. E. (2000). The prefrontal cortex: response selection or maintenance within working memory? Science, 288(5471), 1656–1660. https://10.1126/science.288.5471.1656

Reuter, M., Rosas, H. D., & Fischl, B. (2010). Highly accurate inverse consistent registration: a robust approach. Neuroimage, 53(4), 1181–1196. 10.1016/j.neuroimage.2010.07.020

Saryazdi, R., Bannon, J., Rodrigues, A., Klammer, C., & Chambers, C. G. (2018). Picture perfect: A stimulus set of 225 pairs of matched clipart and photographic images normed by Mechanical Turk and laboratory participants. Behavior Research Methods, 50, 2498–2510.

Satterthwaite, T. D., Elliott, M. A., Gerraty, R. T., Ruparel, K., Loughead, J., Calkins, M. E., … & Wolf, D. H. (2013). An improved framework for confound regression and filtering for control of motion artifact in the preprocessing of resting-state functional connectivity data. Neuroimage, 64, 240–256. 10.1016/j.neuroimage.2012.08.052

Schaul, T., Horgan, D., Gregor, K., & Silver, D. (2015, June). Universal value function approximators. In International conference on machine learning (pp. 1312–1320). PMLR.

Stachenfeld, K. L., Botvinick, M. M., & Gershman, S. J. (2017). The hippocampus as a predictive map. Nature Neuroscience, 20(11), 1643–1653. 10.1038/nn.4650

Tervo, D. G. R., Tenenbaum, J. B., & Gershman, S. J. (2016). Toward the neural implementation of structure learning. Current Opinion in Neurobiology, 37, 99–105. 10.1016/j.conb.2016.01.014

Tomov, M. S., Schulz, E., & Gershman, S. J. (2021). Multi-task reinforcement learning in humans. Nature Human Behaviour, 5(6), 764–773. 10.1038/s41562-020-01035-y

Tustison, N. J., Avants, B. B., Cook, P. A., Zheng, Y., Egan, A., Yushkevich, P. A., & Gee, J. C. (2010). N4ITK: improved N3 bias correction. IEEE Transactions on Medical Imaging, 29(6), 1310–1320. 10.1109/TMI.2010.2046908.

Weiskopf, N., Hutton, C., Josephs, O., & Deichmann, R. (2006). Optimal EPI parameters for reduction of susceptibility-induced BOLD sensitivity losses: a whole-brain analysis at 3 T and 1.5 T. Neuroimage, 33(2), 493–504. 10.1016/j.neuroimage.2006.07.029

Wimmer, G. E., & Büchel, C. (2019). Learning of distant state predictions by the orbitofrontal cortex in humans. Nature Communications, 10(1), 2554. 10.1038/s41467-019-10597-z

Wittkuhn, L., & Schuck, N. W. (2021). Dynamics of fMRI patterns reflect sub-second activation sequences and reveal replay in human visual cortex. Nature Communications, 12(1), 1795. 10.1038/s41467-021-21970-2

Wittkuhn, L., Krippner, L. M., Koch, C., & Schuck, N. W. (2024). Replay in human visual cortex is linked to the formation of successor representations and independent of consciousness. BioRxiv. 10.1101/2022.02.02.478787

Xia, L., & Collins, A. G. (2021). Temporal and state abstractions for efficient learning, transfer, and composition in humans. Psychological Review, 128(4), 643–666. 10.1037/rev0000295

Zhang, Y., Brady, M., & Smith, S. (2001). Segmentation of brain MR images through a hidden Markov random field model and the expectation-maximization algorithm. IEEE Transactions on Medical Imaging, 20(1), 45–57. 10.1109/42.906424.

