## Supplement for "Neural evidence that humans reuse strategies to solve new tasks"

**Supplementary Information**

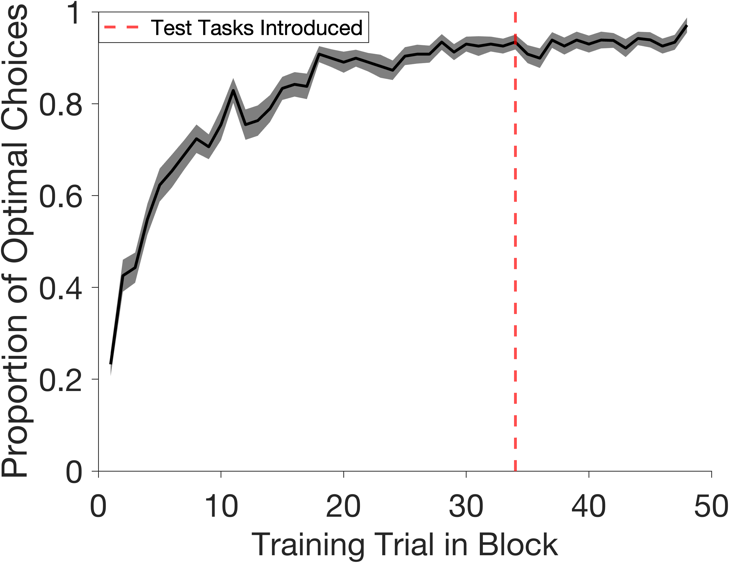

*Figure S1.* Learning dynamics on training tasks. This plot shows the proportion of optimal choices for the 48 training trials per block (i.e. as a function of time). The black line shows the mean across participants and shading indicates the standard error of mean. The red line indicates the point in each block when test tasks are introduced. The earliest this could occur was trial 34. Performance begins around chance (M_trial 1_=0.24, SD_trial 1_=0.16) and reaches an asymptote by the time test tasks are introduced (M_trial 33_=0.93, SD_trial 33_=0.10).

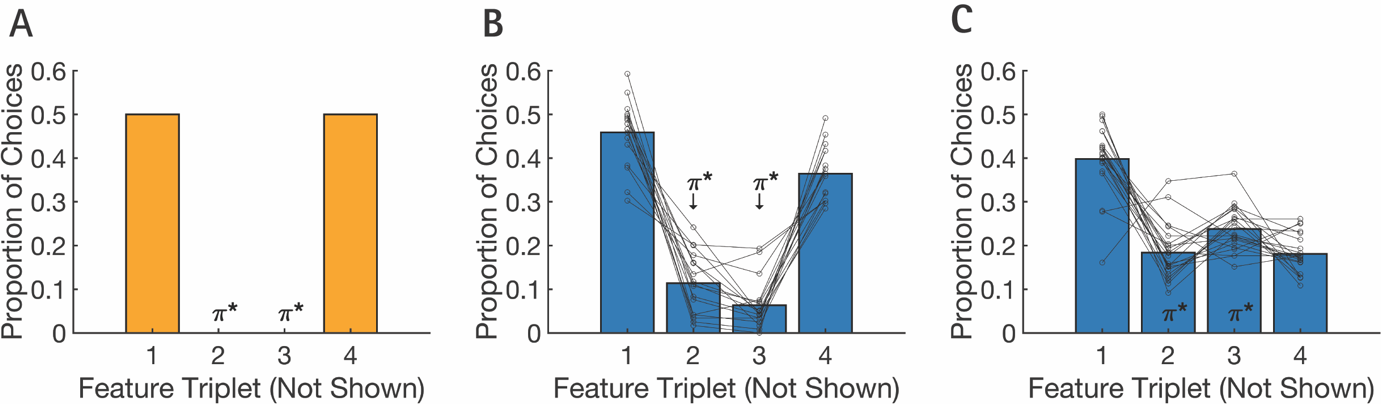

*Figure S2*. Subgroup analysis. Exploratory analysis of participant choices on test tasks revealed two subgroups. Each plot shows the proportion of choices on test tasks (Y-axis) that lead to each feature triplet (X-axis). π* denotes feature triplets associated with optimal test policies. **A:** The predicted choice profile across test tasks according to an SF&GPI algorithm. **B:** The choice profile for a subgroup of participants (n=17) showing a strong recapitulation of the SF&GPI predictions. The choice profile for the remaining participants (n=19) showing a partial recapitulation. **B-C**: Dots show individual participant data. Lines connect individual participant data across the feature triplets.

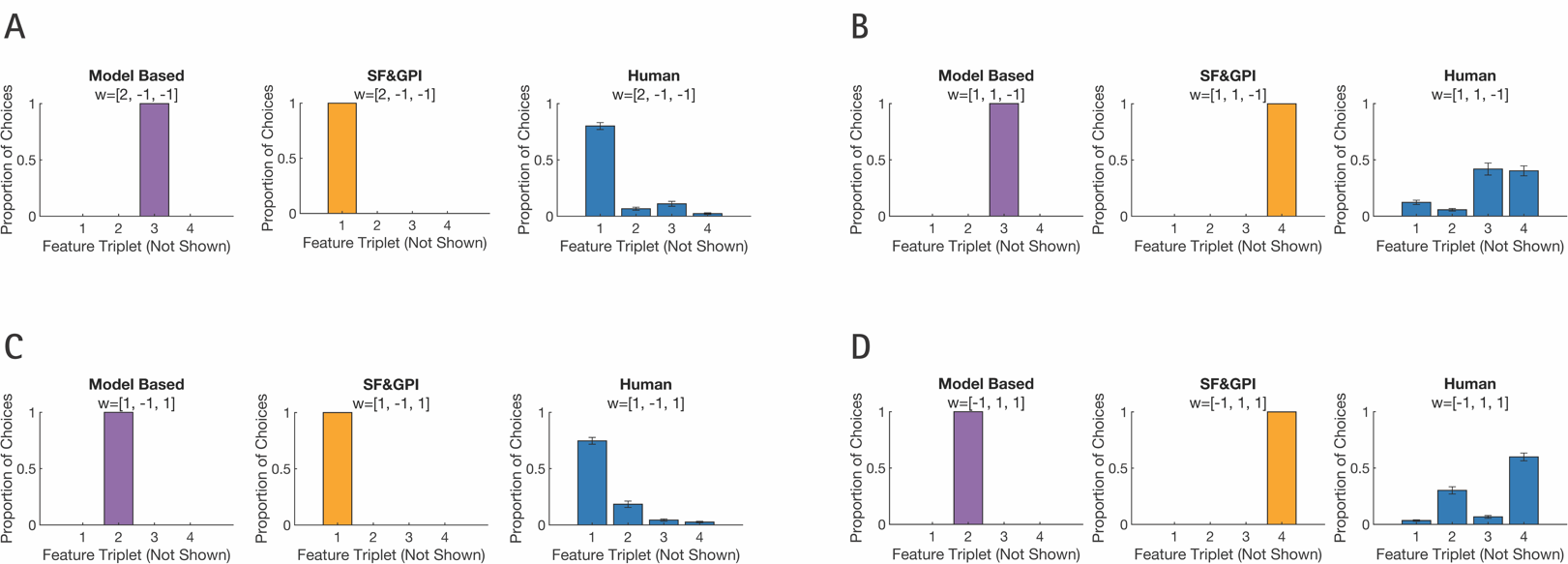

*Figure S3*. Choice behaviour on individual test tasks. Participants’ most common action matched the SF&GPI predictions in 3/4 cases. Each plot shows the proportion of choices (Y-axis) that lead to each feature triplet (X-axis). Each panel shows the choice profile for an individual test task. The cue for an individual task was a weight vector, *w*, which is indicated in the plot titles. These cues were called market values during the gem collector game. Each panel includes three plots: the theoretical predictions of a model-based algorithm, the theoretical predictions of an SF&GPI algorithm and empirical choices from human participants. Bars indicate the standard error of the mean in plots with human data. To formally examine human choice profiles on an individual task level, we compared whether participants made the SF&GPI predicted choice significantly more often than the alternatives. For example, using data from the test task *w*=[2,-1,-1] we compared whether the proportion of choices leading to feature triplet 1 was significantly higher than the proportions leading to feature triplet 2, 3 and 4. Repeating this process for each individual test task revealed that participants made the SF&GPI-predicted choice significantly more often in all cases except one (all *t*-values without the exception > 4.54, all corrected *p*-values without the exception < 0.001). The exception can be seen in panel **B** where the proportion of SF&GPI-predicted choices and MB-predicted choices were comparable (M_ɸ(4)_=40.24% of test trials, SD_ɸ(4)_=27.40, M_ɸ(3)_=41.83%, SD_ɸ(3)_=32.91, *t*(37)=-0.166, corrected *p*=0.869). Choice proportions were compared using paired samples t-tests and *p*-values were corrected for the 12 tests in this section using the Bonferroni-Holm correction.

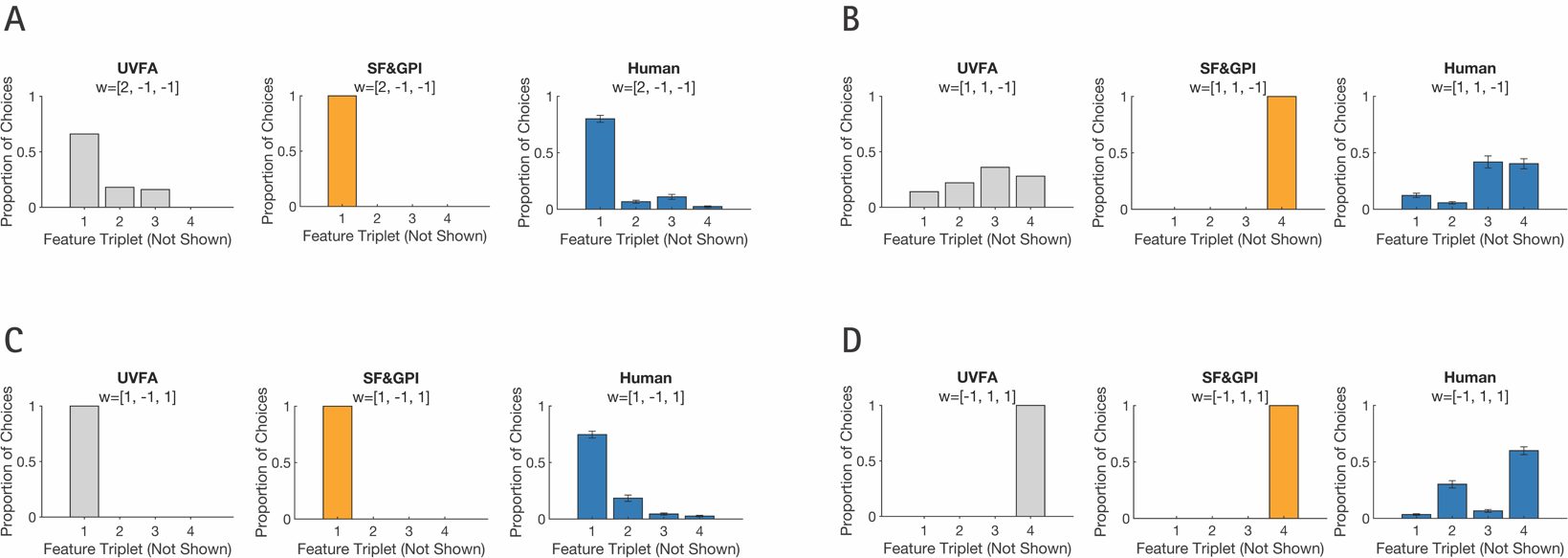

*Figure S4*. Predictions from a Universal Value Function Approximator (UVFA) algorithm. Each plot shows the proportion of choices (Y-axis) that lead to each feature triplet (X-axis). Each panel shows the choice profile for an individual test task. The cue for an individual task was a weight vector, *w*, which is indicated in the plot titles. These cues were called market values during the gem collector game. Each panel includes three plots: the theoretical predictions of a UVFA algorithm, the theoretical predictions of an SF&GPI algorithm and empirical choices from human participants. Bars indicate the standard error of the mean in plots with human data. UVFA predictions were generated post hoc using the implementation in Tomov et al. (2021).

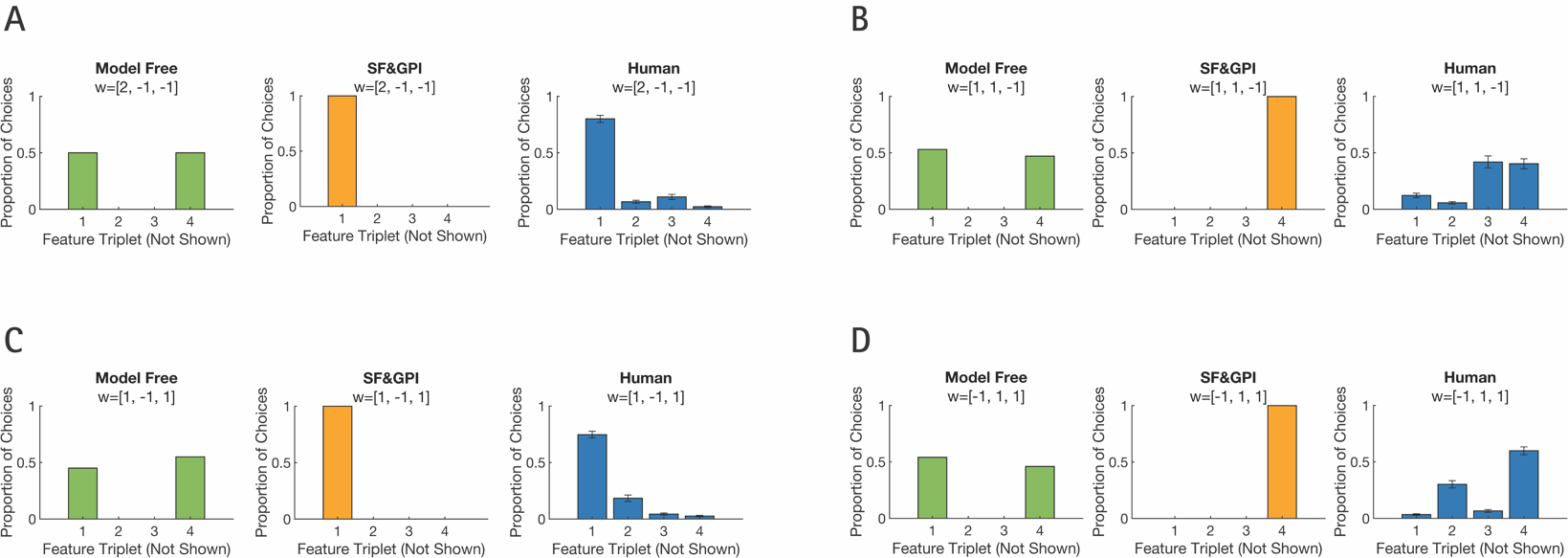

*Figure S5*. Choices on individual test tasks were not explained by model-free perseveration. A model-free algorithm learns the optimal policies on training tasks but has no way to evaluate their success on test tasks. The result is that the algorithm reuses the optimal training policies in an unselective fashion. Each plot shows the proportion of choices (Y-axis) that lead to each feature triplet (X-axis). Each panel shows the choice profile for an individual test task. The cue for an individual task was a weight vector, *w*, which is indicated in the plot titles. These cues were called market values during the gem collector game. Each panel includes three plots: the theoretical predictions of a model-free algorithm, the theoretical predictions of an SF&GPI algorithm and empirical choices from human participants. Bars indicate the standard error of the mean in plots with human data.

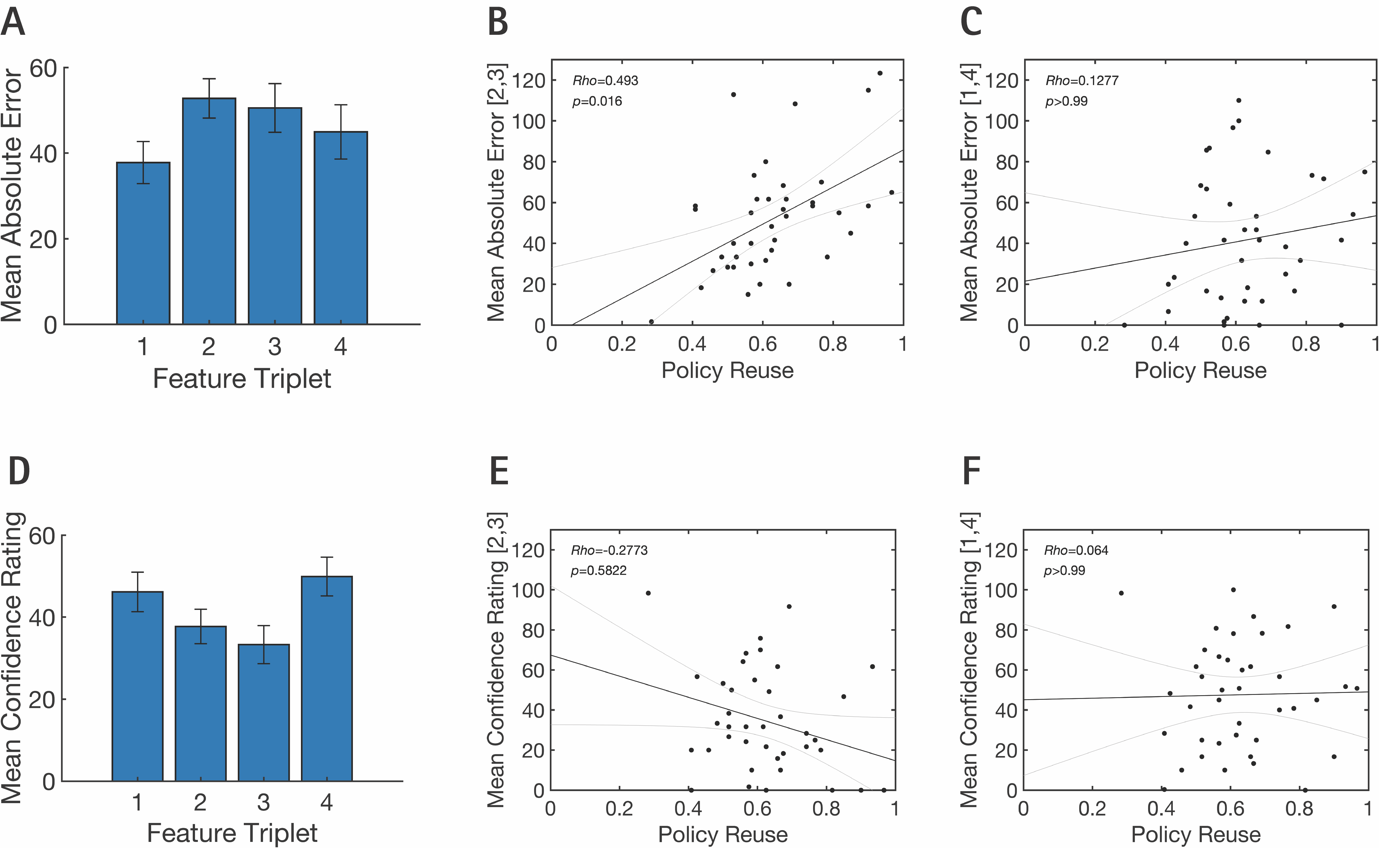

*Figure S6*. Post scan feature estimates and confidence ratings. Participants were asked to estimate the number of triangle, square and circle gems that had appeared in each city during the final block of the scanning task. Estimates were restricted to a range between 0-250 gems. Participants were also asked to rate their confidence for each estimate on a scale from 0 (not confident at all) to 100 (completely confident). **A:** The mean absolute error (Y-axis) for each feature triplet (X-axis). The mean error in each bar is an average across the three estimates provided for each triplet. No significant differences in mean absolute error were detected when correcting for multiple comparisons (all corrected *p* values>0.09). **B:** The relationship between the mean error for feature triplets associated with the suboptimal training policies and policy reuse. Feature triplets 2 and 3 were associated with suboptimal training policies. Policy reuse is the proportion of test trials in which the more rewarding optimal training policy was selected. We detected a significant correlation between these variables after correcting multiple comparisons, suggesting that individuals with less accurate estimates of features linked to the suboptimal training policies were less likely to select those policies on test trials. **C:** The same plot as B, except showing the mean absolute error for feature triplets associated with the optimal training policies. **D:** Mean confidence ratings (Y-axis) for each feature triplet (X-axis). Mean confidence in each bar is an average across the three ratings provided for each triplet. Participants were significantly more confident in their ratings for feature triplets associated with the optimal training policies [ɸ(1) and ɸ(4)] compared to ɸ(3) which was associated with one of the suboptimal training policies (t(37)_ɸ(1) vs. ɸ(3)_=3.08, corrected *p*=0.046; t(37)_ɸ(4) vs. ɸ(3)_=3.91, corrected *p*=0.0054, all remaining corrected *p* values>0.05). **E:** The relationship between participants’ confidence in their estimates for the features associated with suboptimal training policies and the proportion of test trials in which the more rewarding optimal training policy was selected. **F:** The same plot as E, except showing the participants’ confidence in their estimates for the features associated with the optimal training policies. **A, D:** Error bars indicate standard error of the mean. **B, C, E, F:** Black lines indicate linear fits to the data and grey lines indicate 95% confidence intervals of the fits. **A-F:** Significance thresholds were corrected for 14 exploratory tests (5 pairwise comparisons for each bar plot and four correlations). The Bonferroni-Holm correction was used to correct for multiple comparisons.

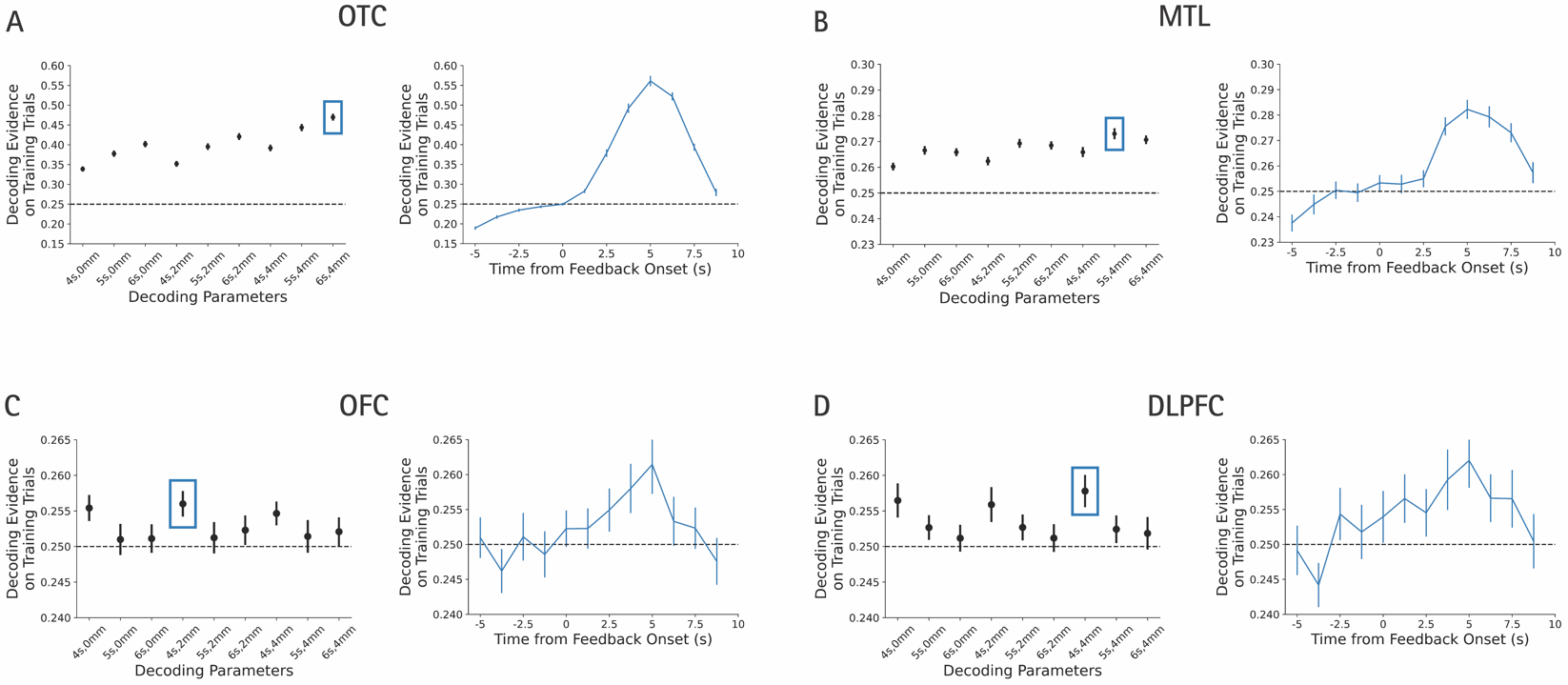

*Figure S7*. Validation of the neural decoding pipeline. To validate our decoding approach, we trained neural decoders to distinguish the city observed during feedback on training trials. We then tested how well we could recover the chosen city on held out training trials. One functional run was held out at a time to assess decoding performance. As training trials were not related to our hypotheses, we used the validation procedure to explore three possible time shifts (+4s, +5s, +6s seconds following feedback onset) and three possible smoothing levels (0mm, 2mm, 4mm) that could be used to optimise decoder training. The validation results were based on 25 iterations. Data were subsampled at random on each iteration to match the number of trials from each class. The highest performing decoding parameters were selected for each ROI based on the validation results. These parameters were later used when testing our hypotheses about neural reactivation during test tasks. **A:** Validation results for occipitotemporal cortex (OTC). The left subplot shows the mean decoding evidence for the selected city on held out training trials for each combination of decoding parameters. The mean is taken across a 5s window (3.75s-8.75s) following feedback onset. Bars indicate standard error of the mean. The blue box denotes the decoding parameters with the highest validation performance. The right subplot shows the decoding time course based on those parameters. **B-D:** The remaining panels follow the same structure as **A** but present validation results for different brain regions. These include the medial temporal lobe (MTL) in **B**, the orbitofrontal cortex (OFC) in **C** and the dorsolateral prefrontal cortex (DLPFC) in **D**.

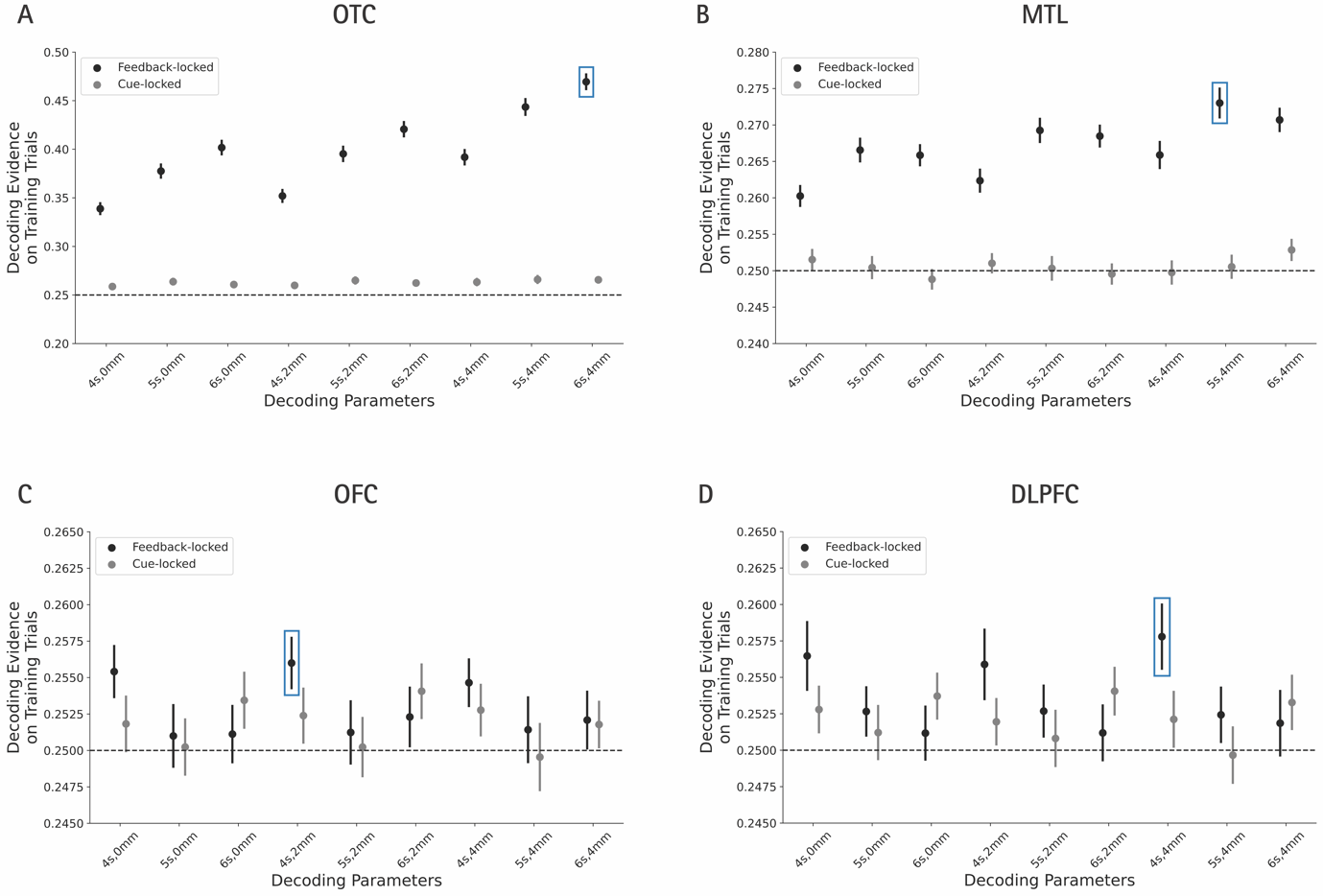

Figure S8. Comparison of cue-locked and feedback-locked decoding. It is possible that the activation of city stimuli could occur directly following task cue onset. We therefore performed additional validation tests, examining the decoding evidence on training tasks when decoder training was locked to task cue onset, rather than feedback onset. Decoding based on feedback onset was found to be more effective in all ROIs. Panels **A-D** were generated using the same approach described in Figure S7. For cue-locked decoding, the decoder was trained using data with a time shift of +4s, +5s, or +6s from cue onset and evaluated from cue onset on held-out trials. Decoding evidence above represents evidence for the city selected on each training trial, averaged over a 5s window (3.75-8.75s) following the relevant event onset (cue presentation or feedback presentation). **A:** Validation results for occipitotemporal cortex (OTC). Bars indicate standard error of the mean. The blue box denotes the decoding parameters with the highest validation performance. **B-D:** The remaining panels follow the same structure as **A** but present validation results for different brain regions. These include the medial temporal lobe (MTL) in **B**, the orbitofrontal cortex (OFC) in **C** and the dorsolateral prefrontal cortex (DLPFC) in **D**.

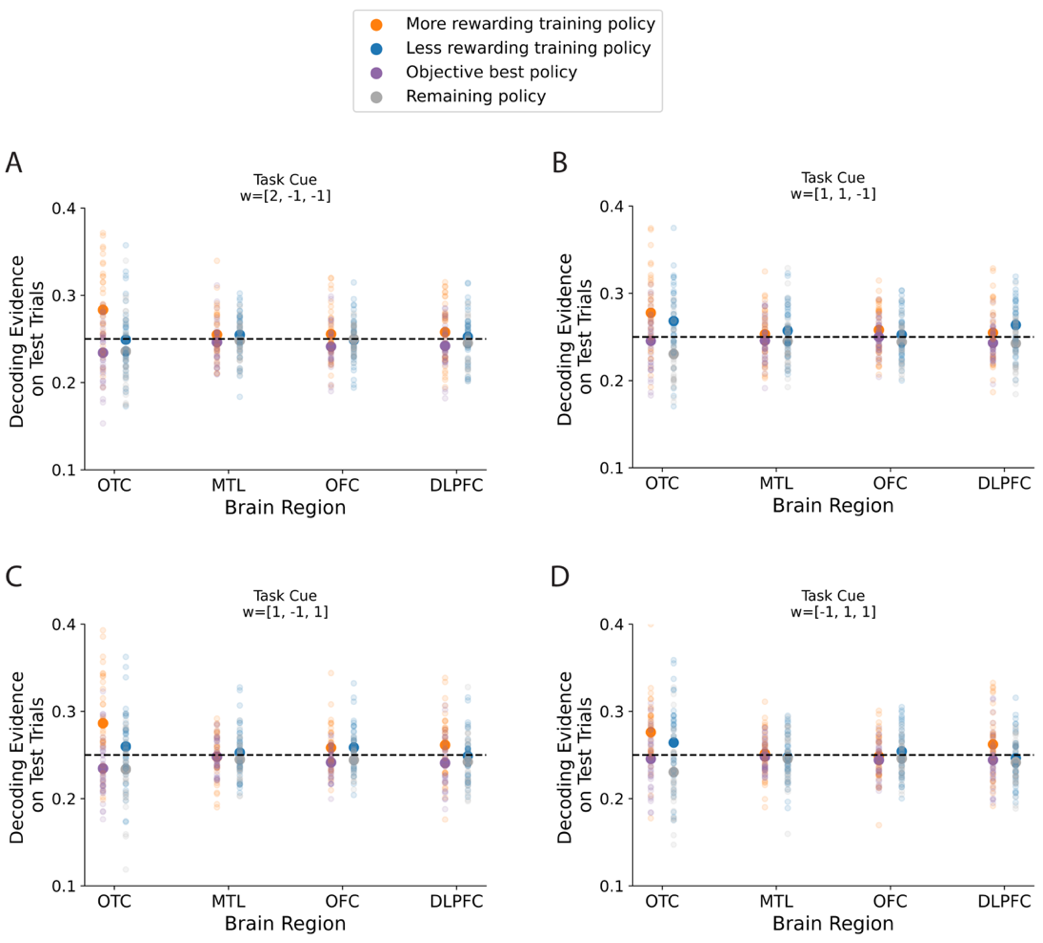

*Figure S9.* Decoding evidence for policies on each individual test task. Each panel shows average decoding evidence during a specific test task (y-axis) as a function of brain region (x-axis). Colours denote different policy categories and small translucent circles show data from individual participants. Panel **B** is especially relevant for testing whether the neural data are more consistent with SF&GPI or Universal Value Function Approximation (UVFA). The theoretical choice profiles for SF&GPI and UVFA are distinct for test task w=[1,1,-1] (Fig. S4). The theoretical choice profile for UVFA predicts: 1) that the more rewarding training policy and objective best policy will be activated on test task w=[1,1,-1] and 2) that the decoding evidence for these two policies will be comparable in magnitude. In contrast, we found that the more rewarding training policy was activated in OTC (*t*(37)=3.87, corrected *p*=0.003) but did not detect evidence that the objective best policy was activated (*t*(37)=-0.90, corrected *p*=0.750). Decoding evidence for the more rewarding training policy was also significantly higher than the objective best policy (*t*(37)=3.21, corrected *p*=0.014). Equivalent tests in DLPFC were not significant (*t*-values<1.66, corrected *p*-values>0.094). These follow up tests were restricted to OTC and DLPFC based on the decoding results in Fig. 3 of the main text. *p*-values were corrected for 6 tests using the Bonferroni-Holm correction (2 policy categories x 2 ROIs and 2 follow up tests assessing the difference in decoding evidence in each ROI).

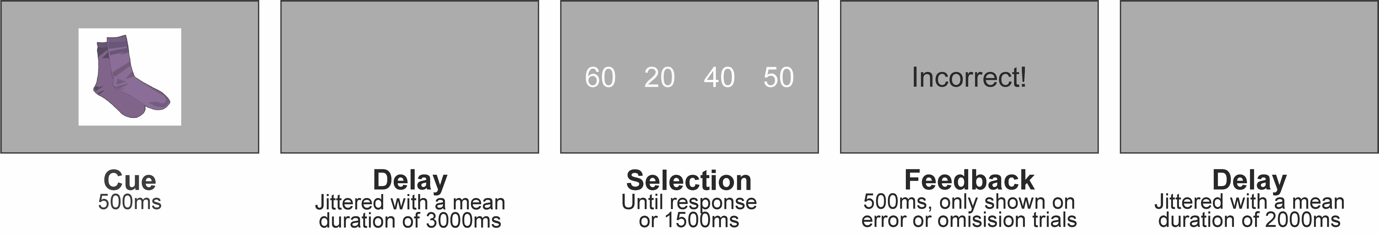

*Figure S10.* Associative memory paradigm used to train the feature decoders. On each trial, participants were shown a retrieval cue and need to select a corresponding target stimulus. Targets were either feature numbers or cities that would later appear during the gem collector game. Participants were pre-trained on the associations before scanning and two retrieval cues were used per target stimulus. Feature decoders were then trained using neural patterns that arose from the response phase onset. For a full description, please see *Session One* and *Feature Decoding* in the Methods.

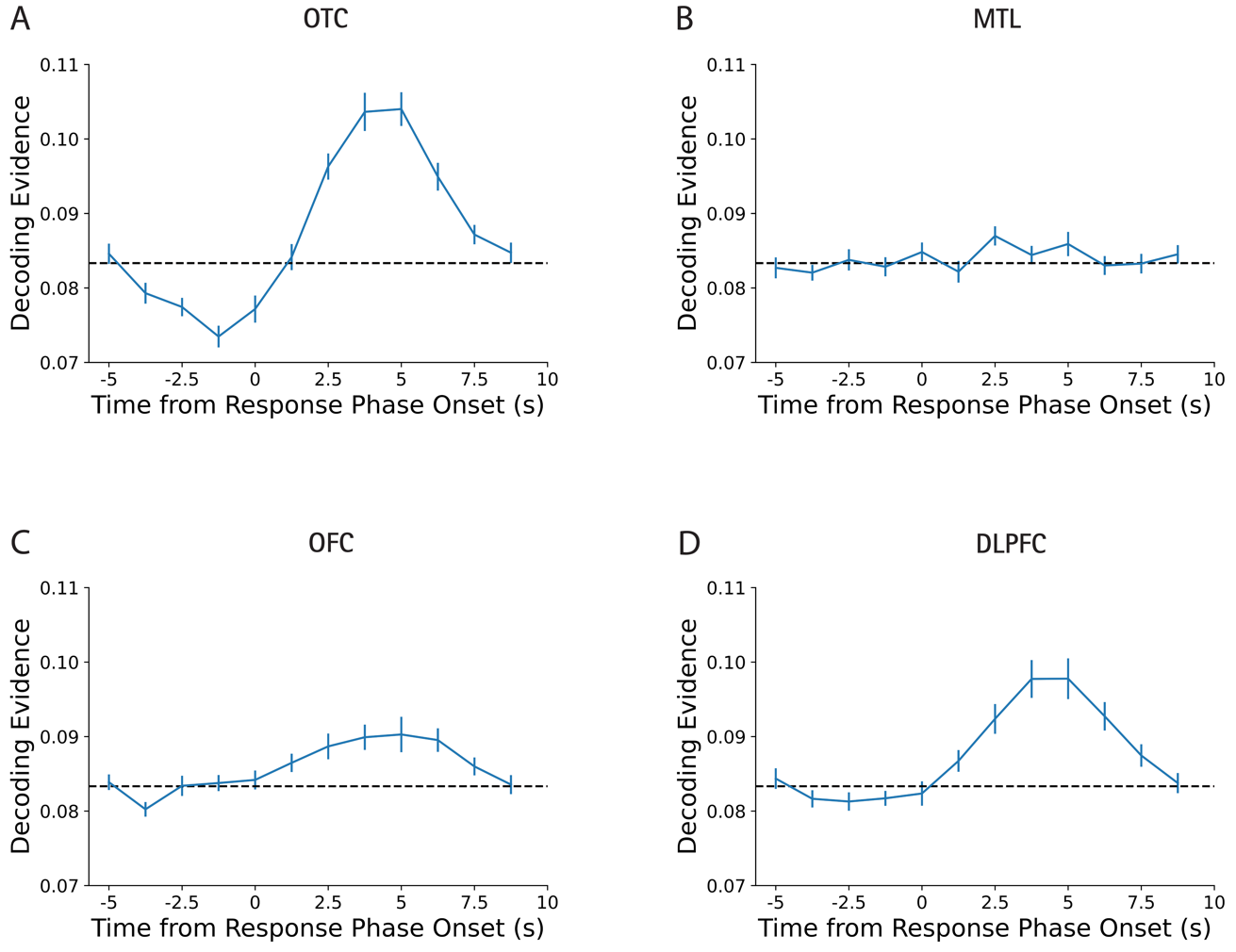

*Figure S11.* Validation tests of feature decoding during the associative memory paradigm. To test the validity of our feature decoding approach, we trained neural decoders to distinguish the target number presented during the response phase. Response phase onset was used for training to be as consistent as possible as the policy decoding approach in Figure S7, given the associative memory paradigm did not have feedback after correct responses. The validation tests used the optimal smoothing and time shift for each ROI identified in Figure S7. Once the feature decoders were trained, we tested how well we could recover the target number on held out trials. One functional run was held out at a time to assess decoding performance. The process was repeated 25 times. Data were subsampled at random on each iteration to match the number of trials from each target class. **A:** Decoding time course for occipitotemporal cortex (OTC). Bars indicate standard error of the mean. The dotted line indicates chance based on 12 possible classes. **B-D:** The remaining panels follow the same structure as **A** but present validation results for different brain regions. These include the medial temporal lobe (MTL) in **B**, the orbitofrontal cortex (OFC) in **C** and the dorsolateral prefrontal cortex (DLPFC) in **D**. The target number could be recovered in OTC, OFC, DLPFC but not MTL using this approach.

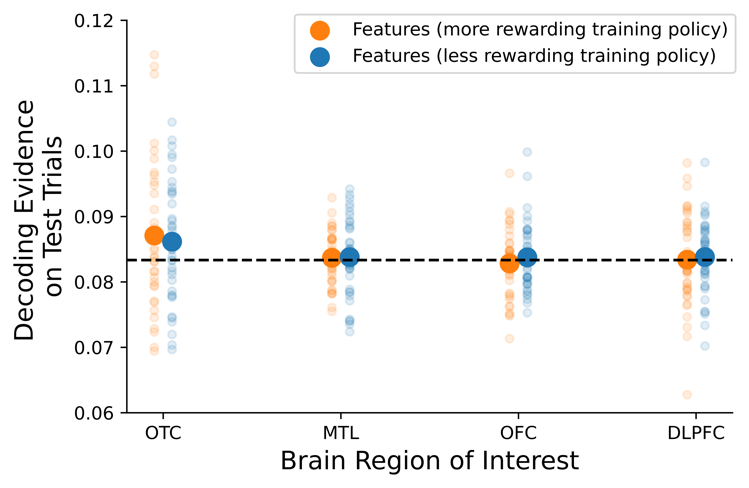

*Figure S12.* Decoding evidence for feature triplets on test tasks. Another neural hypothesis formulated based on the SF&GPI algorithm is that feature triplets associated with the optimal training policies should be reactivated on test trials. This figure shows average decoding evidence for features associated with the more and less rewarding training policies on test trials (y-axis) as a function of brain region (x-axis). Feature information could not be decoded above chance in the four brain regions of interest (corrected *p* values>0.05).

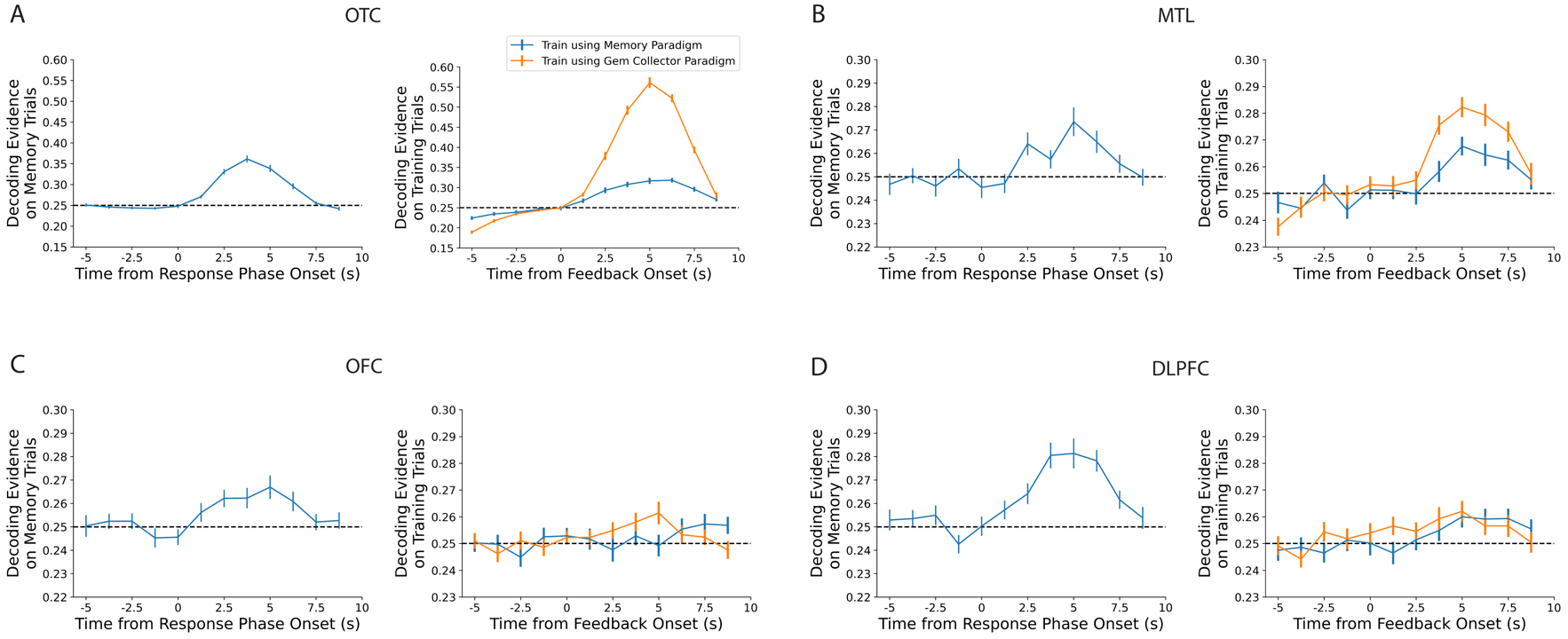

*Figure S13*. Comparison of city decoders trained on session one and session two data. Target cities could be recovered based on neural activity in all ROIs during the associative memory paradigm (session one). However, decoding evidence for cities during the gem collector paradigm (session two) was often higher when decoders were trained on the data from that same paradigm. **A:** Decoding evidence in OTC. The left subplot shows decoding evidence for target cities during the associative memory paradigm. The right subplot shows decoding evidence for cities selected on training trials of the gem collector paradigm, when the decoder was trained on data from session one (blue) or session two (orange). Bars indicate standard error of the mean and the dotted line indicates chance. **B-D:** Panels follow the same structure as **A** but present decoding time courses for different ROIs. **A-D**: Decoder training used the optimal smoothing and time shift for each ROI identified in Figure S7 and 25 subsampling iterations were performed to match the number of trials from each target class. Decoding evidence presented above is cross-validated, based on assessment on held-out test data.
